# Transcriptomic, proteomic and phosphoproteomic underpinnings of daily exercise performance and Zeitgeber activity of endurance training

**DOI:** 10.1101/2020.10.19.345686

**Authors:** Geraldine Maier, Julien Delezie, Pål O. Westermark, Gesa Santos, Danilo Ritz, Christoph Handschin

**Author notes:** Equal first authors (Geraldine Maier, Julien Delezie). Correspondence to:* Christoph Handschin, Biozentrum, University of Basel, Klingelbergstrasse 50/70, CH-4056 Basel, Switzerland.

## Abstract

Timed physical activity might potentiate the health benefits of training. The underlying signaling events triggered by exercise at different times of the day are, however, poorly understood. Here, we found that time-dependent variations in maximal treadmill exercise capacity of naïve mice were associated with energy stores, mostly hepatic glycogen levels. Importantly, running at different times of the day resulted in a vastly different activation of signaling pathways, e.g., related to stress response, vesicular trafficking, repair, and regeneration. Second, voluntary wheel running at the opposite phase of the dark, feeding period surprisingly revealed minimal Zeitgeber (i.e., synchronizing) activity of training. This integrated study provides important insights into the circadian regulation of endurance performance and the control of the circadian clock by exercise. These results are of high importance to understand circadian aspects of training design in athletes and the application of chrono-exercise-based interventions in patients.

**Highlights:**

- Maximal endurance performance is greater in the early morning
- Timed exercise differentially alters the muscle transcriptome and (phospho)-proteome
- Morning exercise triggers energy provisioning and tissue regeneration
- Evening exercise activates stress-related and catabolic pathways
- Training exerts poor Zeitgeber activity on the muscle and liver clocks

## Introduction

Almost all aspects of mammalian physiology undergo changes relative to the time of the day. Many of these variations are, directly or indirectly, driven by the circadian clock, an evolutionary conserved time-keeping mechanism that is present in virtually all cells of the body (Buhr and Takahashi, 2013). Adequate timing of physical activity might have been of particular evolutionary importance to synchronize movement with predator-prey interactions, implying a strong control by the circadian clock, both at the central level to control behavior and in skeletal muscle for adequate functionality (Albrecht and Eichele, 2003). Inversely, adaptation of clock-controlled physiological parameters, for example, anticipation and execution of metabolic pathways important for foraging, optimally can be influenced by activity patterns to adapt to changes in the external environment (Hughes and Piggins, 2012). Thus, in addition to potentially being downstream of clock control, muscle activity has also been proposed as upstream Zeitgeber to synchronize peripheral clocks, and the application of chrono-exercise put forward in various diseases characterized by abnormal circadian rhythmicity (Gabriel and Zierath, 2019; Gutierrez-Monreal et al., 2020).

Consistent with the circadian modulation of muscle physiology, hundreds of transcripts in skeletal muscle oscillate with a 24-h period in humans and mice (McCarthy et al., 2007; Perrin et al., 2018). Moreover, insulin sensitivity, mitochondrial respiration, glucose, and lipid-related metabolites likewise follow similar patterns in muscle tissue (Dyar et al., 2018; Loizides-Mangold et al., 2017; Sato et al., 2018). In line with this, daily variations in resistance and endurance exercise peak performance have been reported during the normal active phase in humans (Mirizio et al., 2020) and rodents (Ezagouri et al., 2019) in some, but not all trials (Knaier et al., 2019; Mirizio et al., 2020). The robustness and timing of such performance peaks seems highly variable, depending on a multitude of parameters including chronotype, time from awakening, muscle and liver glycogen levels, nutritional status, and temperature (Facer-Childs and Brandstaetter, 2015; Hearris et al., 2018). It thus is unknown whether and how the intrinsic muscle clock machinery influences the physiological and molecular responses of skeletal muscle to exercise and ultimately physical performance. Inversely, a potential Zeitgeber activity of training also is unclear. Training studies at different times of the day suffer from confounding aspects such as light-mediated inhibition of voluntary locomotion in animals, or restricted analysis of reporter gene-based approaches. Therefore, whether the enhanced effects of timed exercise training on health parameters in both clinical and preclinical contexts (Savikj et al., 2019; Yamanaka et al., 2015) depend on the Zeitgeber properties of exercise remains to be investigated (Kemler et al., 2020; Sato et al., 2019).

To address open questions about the cross-regulation of exercise and the circadian clock, we first evaluated effects of timed exercise at the systemic and muscle cellular levels by assessing maximal treadmill exercise capacity across the 24-h light-dark (LD) cycle. We furthermore dissected the transcriptome and (phospho-)proteome responses of working muscles at two distinct phases of the LD cycle. Second, to investigate the potential Zeitgeber activity of exercise, we used a skeleton photoperiod (SPP) in combination with restricted wheel-running access to interrogate the consequences of scheduled daytime voluntary training on skeletal muscle gene and protein regulation.

## Results

### Time-of-day-dependent variations in mouse treadmill exercise performance

Differences in low- and moderate-intensity treadmill exercise performance were reported within the active phase of wild-type mice (Ezagouri et al., 2019). It is, however, unclear whether mice show broader variations in maximal exercise performance between the light (i.e., resting/inactive) and the dark (i.e., feeding/active) periods. To investigate how the time of the day affects exercise capacity, we challenged different groups of untrained C57BL/6J mice to an acute bout of high-intensity exercise every 4h across the whole 24-h LD cycle and sacrificed them immediately (Ex+0h) or 3h (Ex+3h) after exhaustion was reached (experimental design and protocol, **Fig. 1A**). We found a significant variation of maximal treadmill running capacity, with a peak and trough of performance at Zeitgeber Time 0 (ZT0; light-onset) and ZT12 (light-offset), respectively (**Fig. 1B, S1A**). Importantly, all mice reached exhaustion and displayed a drastic elevation of blood lactate, yet without significant relation to the time of day (**Fig. 1C, S1B**). Conversely, we observed that blood glucose levels significantly dropped at ZT12 and ZT16 upon exercise (**Fig. 1D, S1C**). Lastly, higher serum corticosterone levels were observed in all exercised groups regardless of time (**Fig. S1D**).

**FIGURE 1.**
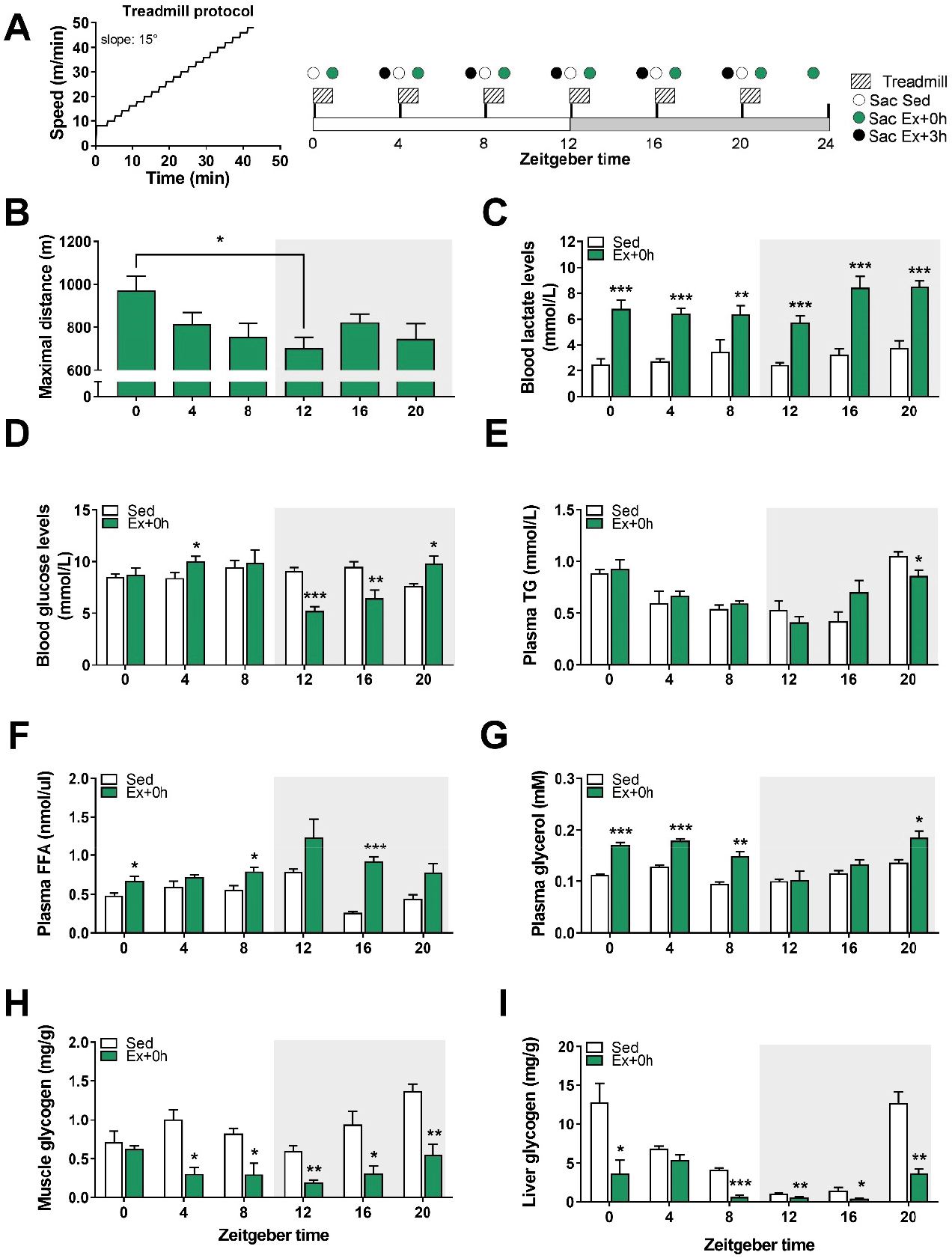
Time-of-day-dependent variations in mouse treadmill exercise performance. (A) Treadmill protocol and experimental scheme: mice were divided into two groups, sedentary (Sed) and exercise (Ex). The latter group was further divided into two groups: sacrificed immediately (+0h) or three hours after exhaustion (+3h). (B) Maximal distance reached at exhaustion. (C) Blood lactate and (D) glucose levels, (E) plasma triglyceride (TG), (F) free fatty acid (FAA), and (G) glycerol levels. (H) Muscle and (I) liver glycogen levels. Data is shown as the average ± SEM (n=3 in Sed and Ex+0h groups). * P < 0.05; ** P < 0.01; *** P < 0.001. One-way ANOVA (B) and Unpaired Student’s t-test (C-I).

To further evaluate the metabolic outcome of maximal treadmill exercise across the day, we measured circulating energy substrates, muscle, and liver glycogen levels immediately after exercise (Ex+0h). We observed that plasma triglyceride (TG) levels were mainly unchanged at exhaustion (**Fig. 1E**), which is consistent with circulating TG not being the primary source of energy during moderate to high-intensity exercise in untrained animals (Hargreaves and Spriet, 2018). Conversely, plasma free fatty acids (FFA) and glycerol levels, a lipolytic marker, were affected by treadmill exercise in a time-dependent manner (**Fig. 1F, G**). Lastly, muscle glycogen stores were consistently reduced by exercise, except for ZT0, the time at which mice show their greatest performance (**Fig. 1H**). In contrast, hepatic glycogen stores, an essential source of glucose during exercise (Richter and Hargreaves, 2013), were significantly impacted by exercise across the LD cycle except for ZT4 (**Fig. 1I**).

Altogether, these data demonstrate that mice are surprisingly better at performing a maximal running test in the early light phase of the 24-h LD cycle. Furthermore, we show that when basal hepatic glycogen stores are low (i.e., in the early dark phase), mice are unable to sustain a prolonged workout and to maintain homeostatic blood glucose levels.

### Exercise around the clock induces broad and time-dependent gene responses in skeletal muscle

Using qPCR, we next evaluated the expression of genes that are part of the immediate, early response of skeletal muscle to exercise across the day (see method for details about the statistical analysis and **Fig. S1E**; for each transcript, normalized data are provided as bar graphs, while raw data are provided as daily line graphs). We found that nuclear receptor subfamily 4 group A member 3 (*Nr4a3*) and activating transcription factor 3 (*Atf3*), both exercise-responsive genes (Fernandez-Verdejo et al., 2017; Pillon et al., 2020), were induced immediately after exercise and remained elevated 3h post-exercise (**Fig. 2A, S2A**). Conversely, while the expression of the peroxisome proliferator-activated receptor γ coactivator 1α 4 (*Ppargc-1α4*) isoform induced by resistance exercise (Ruas et al., 2012) was up-regulated 3h post-exercise (**Fig. S2A**), the *Ppargc-1α1* isoform particularly involved in oxidative muscle remodeling (Lin et al., 2002) was mainly induced directly after exercise (**Fig. S2A**). Moreover, interleukin-6 (*Il-6*), a myokine implicated in muscle glycolysis and adipose fat lipolysis (Pedersen and Febbraio, 2008), was increased immediately post-exercise, especially in the late part of the day and at night (i.e., from ZT8-20) (**Fig. 2A**). Finally, the vascular endothelial growth factor A (*Vegfa*), an important regulator of muscle regeneration, exercise adaptation, and angiogenesis (Delavar et al., 2014), was only induced when exercise was performed during daytime (**Fig. 2A**).

**FIGURE 2.**
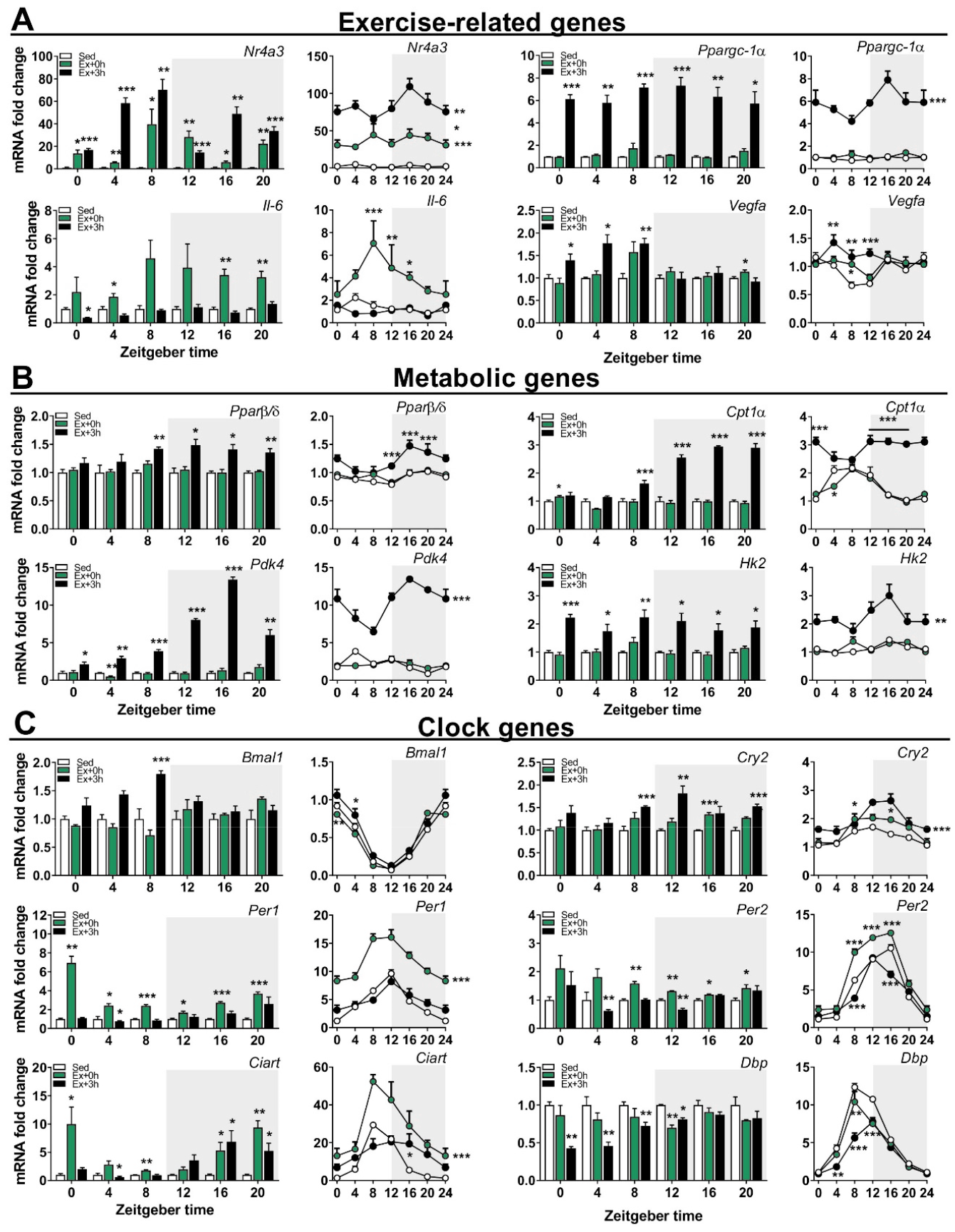
Exercise around the clock induces broad and time-dependent gene responses in skeletal muscle. Gene expression in sedentary (Sed) and exercised mice at (Ex+0h) and 3h (Ex+3h) after exhaustion. (A) Exercise-related genes, (B) Metabolic genes, (C) Clock genes categories. Expression values were determined by qPCR and normalized to *Hprt*. Data in bar graph is shown as the average fold-change ± SEM (n = 3) relative to the expression in Sed set to 1 (see method for details on normalization). Data in line graph is shown as the average fold-change ± SEM (n = 3) relative to the expression in the Sed ZT0 group set to 1. * P < 0.05; ** P < 0.01; *** P < 0.001. Unpaired Student’s t-test (Bar graphs) and One-way ANOVA (Line graphs).

We likewise measured genes involved in the regulation of lipid and glucose metabolism: some were induced at the day-night transition (e.g., peroxisome proliferator-activated receptor β/δ [*Pparβ/δ*]; **Fig. 2B**), induced at night (e.g., the liver-specific isoform Carnitine palmitoyltransferase-1α [*Cpt1α*] and pyruvate dehydrogenase kinase 4 [*Pdk4*]; **Fig. 2B**), broadly induced, (e.g., hexokinase 2 [*Hk2*]; **Fig. 2B**), or unchanged by exercise (e.g., the muscle-specific isoform [*Cpt1β*]; **Fig. S2B**).

Finally, we assessed the expression of core circadian clock genes. We observed changes in the expression of brain and muscle arnt-like (*Bmal*)*1*, circadian locomotor output cycles kaput (*Clock*), cryptochrome *(Cry)1, Cry2*, period *(Per)1, Per2, Per3*, circadian associated repressor of transcription (*Ciart*), retinoic acid receptor-related receptor α (Rorα), *Rorγ*, nuclear receptor subfamily 1 group D member 1 (*Nr1d1, Rev-Erba), Nr1d2 (Rev-Erbβ*) and albumin D box-binding protein (*Dbp*), while only Rorβ transcript levels remained unaffected (**Fig. 2C, S2C**). The transcription of most of these clock genes was positively affected by exercise, with some exhibiting a bimodal regulation, e.g., a repression following an induction for *Per1, Per2* or *Ciart* at specific ZTs (**Fig. 2C**). Overall, these data demonstrate that the regulation of prototypical exercise response and metabolic genes is only in part influenced by the time of the day, which for example seems completely irrelevant for *Ppargc-1α.* Moreover, acute endurance exercise bouts extensively affect the expression of core clock genes in the immediate response after fatigue is reached.

### Daytime vs. nighttime treadmill exercise elicits distinct gene signatures in skeletal muscle

In light of our qPCR results, we further explored the transcriptional signatures of working muscles at distinct phases of the LD cycle. We performed RNA-sequencing (RNA-seq) gene expression profiling of muscles harvested at times when treadmill exercise performance differed the most, namely ZT0 (i.e., light onset) and ZT12 (i.e., light offset).

The overall extent of the transcriptional response shifted from the immediate time point (Ex+0) at the early daytime exercise to the late time point (Ex+3) at the early nighttime exercise (**Fig. 3A, B, S3**). Surprisingly, only about 25% of the differentially expressed genes (DEGs; FDR ≤ 0.05) were shared between ZT0 and ZT12, regardless of the time of sacrifice (i.e., Ex+0h vs. Ex+3h). Examples representing this “core program” are the MAF transcription factors *Maff* and *Mafk*, implicated in cellular stress response and detoxification (Katsuoka et al., 2005); the metallothionein1/2 (*Mt1/2*), involved in oxidative stress protection and regulation of hypertrophy via the Akt pathway (Di Foggia et al., 2014); (**Fig. 3C, D**). The expression of most genes, however, differs qualitatively or at least temporally between the two ZT. For instance, in line with the qPCR data, *Per1* and *Ciart* transcripts were significantly induced immediately after exercise when performed at ZT0, but not at ZT12 (**Fig. 3C**).

**FIGURE 3.**
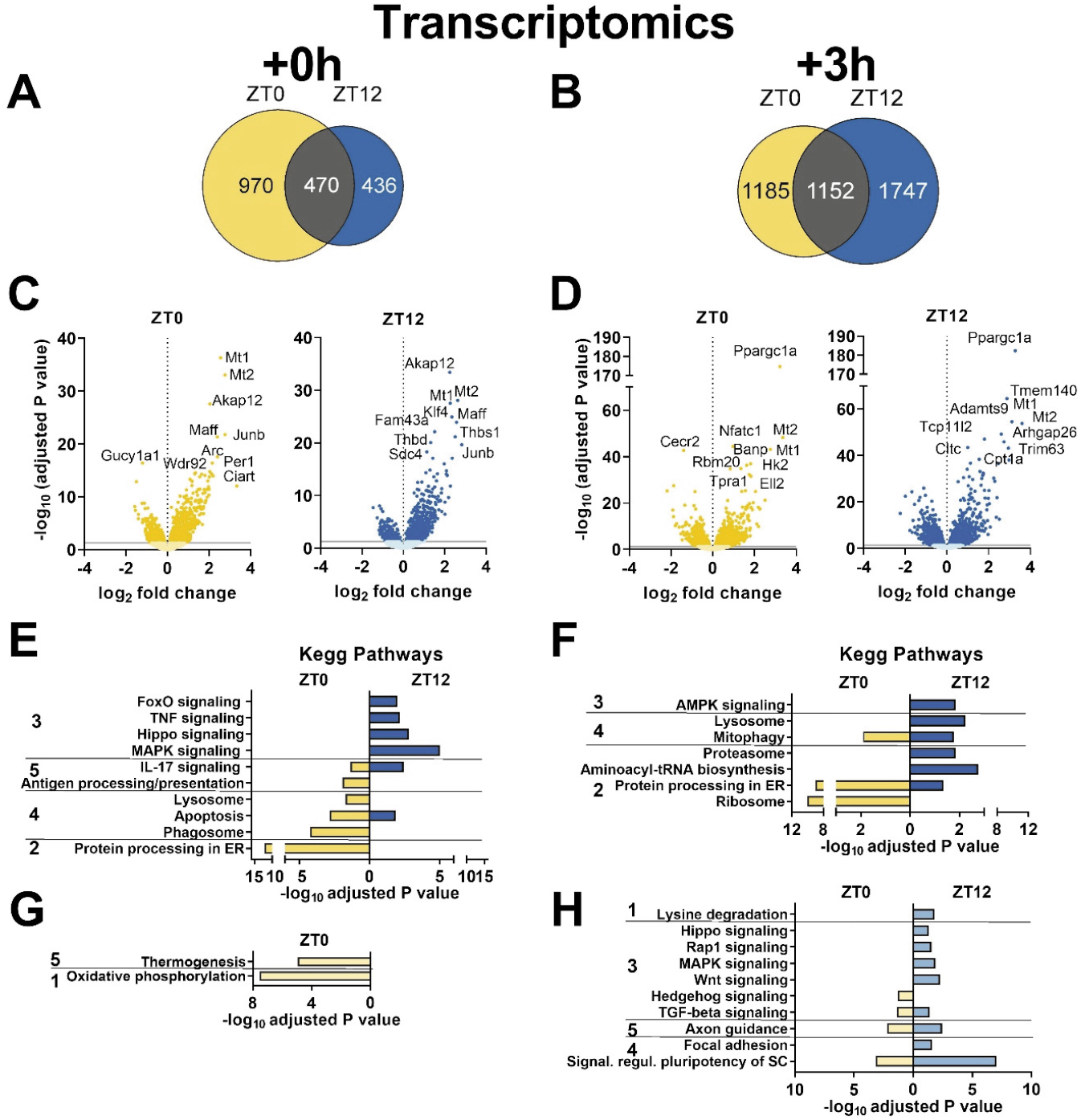
Daytime vs. nighttime treadmill exercise elicit distinct transcriptomic signatures. (A, B) Venn diagrams displaying the number of DEGs immediately (A; Ex+0h) and 3h (B; Ex+3h) after early daytime (ZT0; yellow) or early nighttime (ZT12; blue) exercise, and resulting overlap (gray). (C, D) Volcano plots displaying DEGs (with top 10 indicated) as described above. KEGG analysis (top 5 terms) of (E, F) up-regulated and (G, H) down-regulated genes by exercise as above. KEGG pathway categories: 1. Metabolism; 2. Genetic Information Processing; 3. Environmental Information Processing; 4. Cellular Processes; 5. Organismal Systems.

KEGG pathway enrichment analysis using g:Profiler (Reimand et al., 2016) revealed that exercise at both ZT activated mitophagy, inflammation, and apoptotic pathways (**Fig. 3E, F**). Conversely, exercise at early daytime is linked with a robust immediate increase in genes linked to protein processing, lysosome, and phagosome, associated with a down-regulation of oxidative phosphorylation (**Fig. 3G**), followed by a further increase in protein processing and ribosome-related genes 3h later. In contrast, early nighttime exercise triggers a broad immediate induction in genes related to MAPK and other stress, muscle wasting pathways (i.e., Hippo and FoxO), with an ensuing activation of catabolic signals linked to AMPK, lysosome and proteasome 3h later. At this time point, some of the genes related to the stress signaling pathways that were activated earlier are mostly down-regulated (**Fig. 3H**).

Taken together, the time of the day markedly affects the transcriptional response to an acute bout of endurance exercise. Moreover, broadly speaking, daytime exercise positively regulates transcriptional processes associated with protein synthesis and stability, while nighttime exercise triggers responses associated with energy stress.

### Proteome and secretome changes associated with daytime vs. nighttime treadmill exercise

To investigate whether the exercise-induced transcriptional changes were accompanied by significant modifications at the protein level, we performed mass spectrometric analyses of muscles from mouse groups exercised at ZT0 and ZT12, and identified a total of about 5300 proteins using MaxQuant, specifying a false discovery rate of 5% at the peptide and protein level.

Similar to the transcriptomic data, only a relatively small overlap between the two ZT regarding the levels of differentially affected proteins was observed at either time point (**Fig. 4A-D, S3A**). The control of specific programs was underlined by the small overlap in KEGG terms between ZT0 and ZT12 (**Fig. 4E-4H**), indicating differential regulation of protein expression and pathway activation between time points. For example, the up-regulation of soluble NSF attachment protein receptors (SNARE) interactions in vesicular transport at exhaustion following ZT0 exercise indicates the increase in a class of membrane-associated proteins, which, besides their involvement in neurotransmitter release (Dunant and Israel, 2000; Kasai et al., 2012), regulate GLUT4-containing vesicle trafficking (Cheatham, 2000). In line, the vesicle-associated membrane protein 3 and 8 (VAMP3, 8), syntaxin 6 and 8 (STX6, 8), synaptobrevin homolog YKT6 (YKT6), and of the synaptosomal-associated protein 23 (SNAP23), which are all key mediators of GLUT4 translocation to the cell surface were elevated at this timepoint (Bryant and Gould, 2011; Morris et al., 2020; Zong et al., 2011). Moreover, the Rho family GTPase RAC1, an essential regulator of glucose transport in contracting muscles (Sylow et al., 2017), its upstream regulator Protein tyrosine phosphatase α (PTPα; PTPRA) (Sun et al., 2012), together with the GTPase Ras-related protein (RALA), a known mediator of insulin-dependent glucose uptake (Takenaka et al., 2015), were all consistently elevated. Finally, we observed increased expression of the Rho GTPases RHOA, B, and C, in favor of enhanced GLUT4’s intracellular trafficking (Duong and Chun, 2019). Collectively, these changes imply an activation of glucose uptake into skeletal muscle that seems more specific to the adaptation in early daytime exercise. Moreover, early daytime treadmill robustly initiated the “Complement system” (**Fig. 4E**). Complement activation triggers tissue regeneration in response to muscle injury and inflammation (Zhang et al., 2017). Accordingly, the complement C3a, C4b, C8a, C8b, C9, CFD, and CFB proteins, were all up-regulated. In contrast, the levels of proteins linked to ribosome function were only elevated in early nighttime exercise at 0h, followed by a subsequent decrease at 3h, associated with a strong induction in proteins involved in nicotinamide and nitrogen metabolism (**Fig. 4F, H**). Hence, proteostasis could be more affected at this time point compared to early daytime running.

**FIGURE 4.**
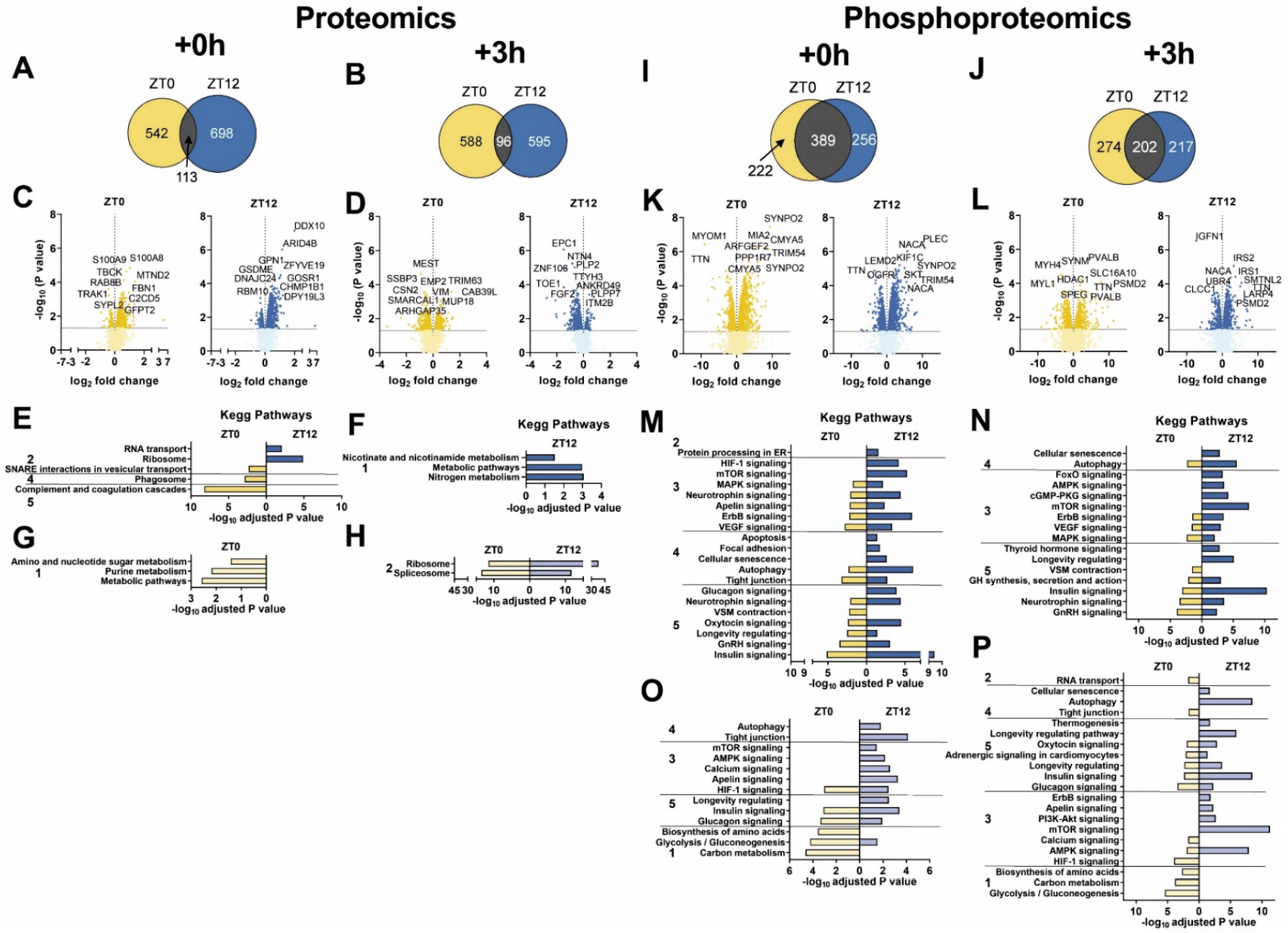
Phospho-/proteomic analyses reveal time-dependent differences in muscle response to early day vs. early nighttime exercise. (A, B) Venn diagrams displaying the number of differentially regulated proteins immediately (A; Ex+0h) and 3h (B; Ex+3h) after early daytime (ZT0; yellow) or early nighttime (ZT12; blue) exercise, and resulting overlap (gray). (C, D) Volcano plots displaying the differentially regulated proteins (with top 10 indicated) as above. KEGG analysis of (E, F) up-regulated and (G, H) down-regulated proteins by exercise as described above. (I, J) Venn diagrams and (K, L) volcano plots displaying the differentially phosphorylated proteins in the same groups as above. KEGG analysis (top 5) for (M, N) increased and (O, P) decreased protein phosphorylation in Ex+0h and Ex+3h groups. KEGG pathway categories: 1. Metabolism; 2. Genetic Information Processing; 3. Environmental Information Processing; 4. Cellular Processes; 5. Organismal Systems.

Skeletal muscle tissue is an endocrine organ, releasing small molecules, so-called myokines, in response to exercise (Delezie and Handschin, 2018). We thus used predictive tools to identify putatively secreted proteins that could influence muscle performance and metabolism at distinct phases of the LD cycle. Of the top 100 proteins exclusively induced in the immediate response to daytime exercise (Ex+0h), we retrieved 25 candidates as potential secreted proteins using SignalP 5.0 and 12 additional using SecretomeP 2.0 (Bendtsen et al., 2004; Nielsen, 2017). For instance, the major urinary proteins (MUP) 3, 17, and 18 were all up-regulated at ZT0 and could potentially be associated with the regulation of systemic blood glucose, as well as liver and skeletal muscle metabolism (Zhou et al., 2009). We, moreover, identified fibrillin-1 (FBN1, also called asprosin), a glucogenic protein hormone that is secreted by adipose cells and recruited at the surface of hepatocytes to increase plasma glucose level (Romere et al., 2016). Nighttime exercise resulted in a smaller number of potentially secreted proteins using the same analysis tools. The 7 predicted proteins included the SPARC-like protein 1 (SPARCL1), a member of the SPARC family of proteins that also includes SPARC/osteonectin, an exercise-regulated myokine with a potential effect on myogenic differentiation (Lee and Jun, 2019).

Overall, these proteomic results highlight the robust activation of time-dependent cellular responses; enhancing glucose metabolism in the early daytime (supported by the predicted secretome) and, conversely, alterations in protein homeostasis when exercise is performed in the early night.

### The phosphorylome of daytime vs. nighttime working muscles

Many proteins are regulated by phosphorylation independently of their expression (Huttlin et al., 2010), in particular upon exercise (Hoffman et al., 2015). To identify signaling pathways that are modulated by exercise at different times of the day, we performed a comprehensive phosphoproteomics analysis, in which approximately 7000 sequence ions carrying post-translational modifications in response to exercise mapping to about 1600 unique proteins were identified (**Fig. 4I-L, S3**).

A larger enrichment of concurrent phosphorylation and dephosphorylation of proteins in stress-related and catabolic pathways was detected in nighttime exercise, both at Ex+0h and Ex+3h (**Fig. 4M-P**). While the α2-subunit of AMPK, the predominant form in the muscle (Garcia and Shaw, 2017), was consistently dephosphorylated on Ser377 by exercise regardless of ZT, only exercise at night induced further changes in the phosphorylation status of both α1/α2 AMPK subunits. Nighttime exercise rapidly phosphorylated α1/α2 at Ser496/491, and we found increased phosphorylation of the critical regulator of autophagy ULK1 on Ser637, which is dependent on AMPK (Mack et al., 2012). The mTOR protein was similarly phosphorylated on Ser1162/1261 upon exercise at both ZT, yet both TSC1 and TSC2 proteins, important integrators of different signaling pathways to control mTOR signaling, were predominantly regulated by nighttime exercise. Emblematic of the activation of the “mTOR pathway”, there were robust posttranslational changes of, e.g., lipin1 (LPIN1), eukaryotic translation initiation factor 4 (EIF4) B, EIF4E binding protein, and ribosomal protein S6 (RPS6). Similarly, key components of the “HIF-1 signaling” pathway, e.g., the glycolytic enzymes enolase 1 (ENO1), phosphofructokinase (PFK) L-M, and aldolase A (ALDOA), displayed unique and/or distinct phosphorylation profiles at night (**Fig. 4M**). In line with the decrease in muscle glycogen content upon exercise at ZT12, the muscle-specific isoform of the glycogen phosphorylase (PYGM), a key enzyme in the first step of glycogenolysis, was specifically phosphorylated on Ser2 and on Ser15, the latter residue is particularly known to enhance phosphorylase activity and the degradation of glycogen (Johnson, 1992). In contrast, these two sites were not phosphorylated in response to early daytime exercise, consistent with muscle glycogen sparing (**Fig. 1H**). Moreover, daytime exercise exclusively enhanced phosphorylation on Ser3/616 residues of the TBC1 domain family member 4 (TBC1D4 or AS160), a strong regulator of GLUT4 trafficking in skeletal muscle (Cartee, 2015; Sakamoto and Holman, 2008). Finally, besides the increased abundance of asprosin in daytime contracting muscles, we observed active posttranslational modifications on Ser2566 and Ser2711, which could warrant a secretory processing to boost hepatic glucose production. We furthermore detected posttranslational modifications of hormone receptors that play a key role in energy metabolism, e.g., the phosphorylation of the glucocorticoid receptor (*Nr3c1*) on T152 and S275 residues only at nighttime, even though plasma corticosterone levels were similarly elevated by exercise at ZT0 and ZT12 (**Fig. S1D**).

Broad changes in the phosphorylation of proteins associated with modulation of muscle contractile properties (i.e., titin; myosin heavy chain 3, 4, and 9; sarcoplasmic reticulum calcium-ATPase 1 and 2) were similarly observed independent of the time of the day (**Fig. 4K, L**). However, the “Vascular smooth muscle contraction” KEGG pathways were exclusively enriched upon daytime exercise, particularly at exhaustion (Ex+0h), indicating a higher engagement of calcium and cAMP signaling pathways. In line, phosphorylation of the calcium voltage-gated channel subunit α 1 S (CACNA1S) protein, a regulator of contractile force in response to stress, fear, and exercise; the so-called “fight-or-flight” response (Catterall, 2015; Emrick et al., 2010), was specific to residue Ser5 and Ser1617/1640 and T700. These sites are not yet characterized but could indicate an up-regulation of calcium channel activity, leading to increased force production. In support of this, expression of the calcium release-activated calcium channel protein 1 (ORAI1), a crucial calcium regulator limiting muscle fatigue (Wei-Lapierre et al., 2013), was exclusively up-regulated in daytime exercised muscles.

Taken together, the phosphoproteomics data mirror the transcriptomics and proteomics inasmuch clear distinctions between different time points were observed, mainly in regard to stress and catabolism. It thus is clear that besides a robust core exercise program, the time of the day has a major impact on the gene expression, protein levels, and posttranslational modifications triggered by acute physical activity.

### Scheduled daytime wheel-running activity in mice exposed to a skeleton photoperiod

While the studies so far pertained to the consequences of the time of the day on the acute exercise response, we next assessed how training at different times of the day affects the clock. Due to the masking (i.e., suppressive) effect of light on physical activity and other nocturnal behavior of mice, voluntary use of the running wheel under normal conditions exclusively spreads throughout the dark period, with a peak in the early hours of the night, with intermittent multiple eating events (Yasumoto et al., 2015). These and other limitations have precluded a comprehensive analysis of prolonged training as potential Zeitgeber under physiological conditions so far. We, therefore, tested the use of a skeleton photoperiod (SPP) in combination with time-restricted wheel and food access to evaluate whether mice would spontaneously run in a wheel during their normal resting phase (see method and **Fig. 5A**). SPPs maintain central clock oscillation (Oishi et al., 2002), reducing the strong inhibition of light on physical activity in mice (Delezie et al., 2016).

**FIGURE 5.**
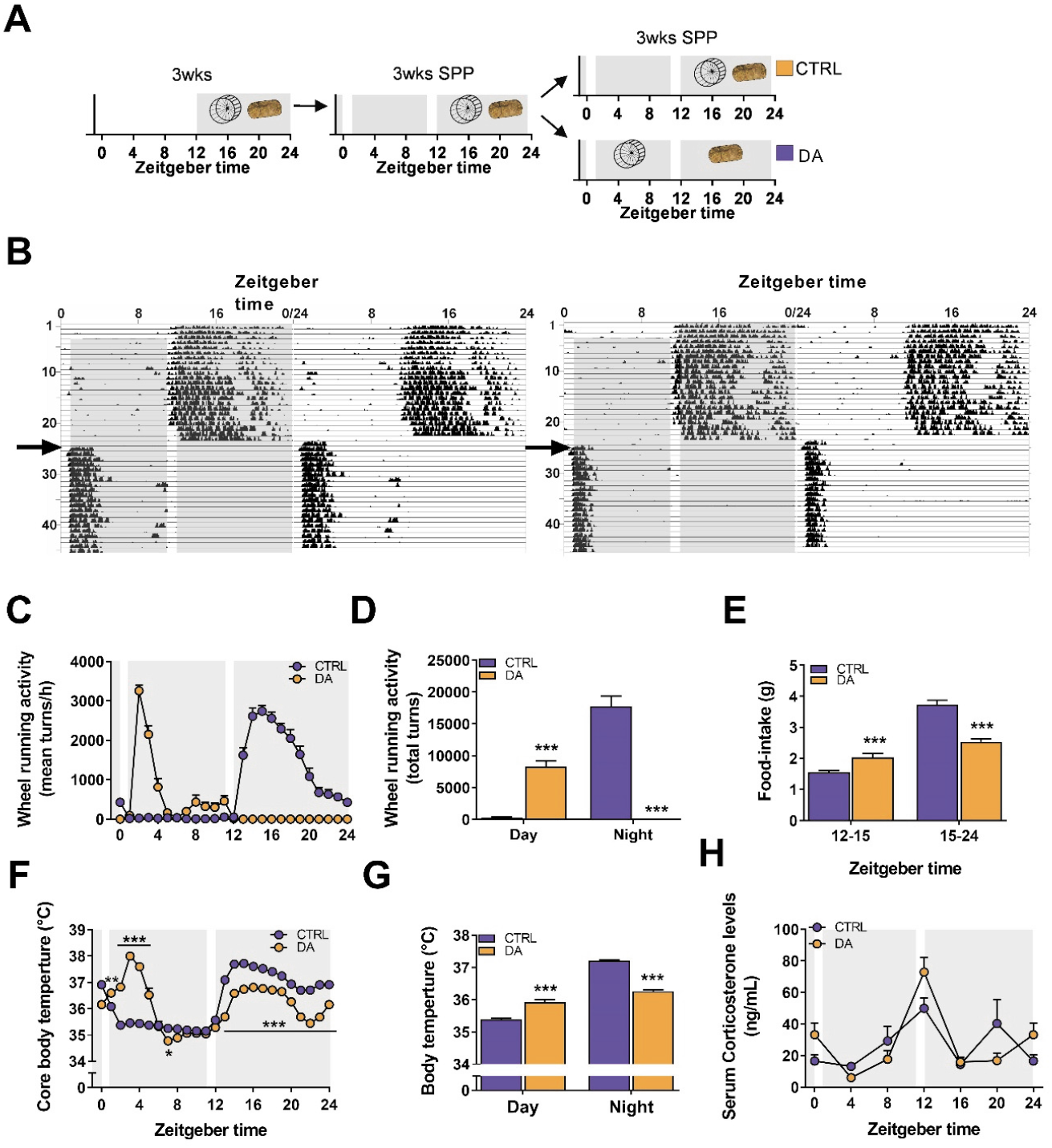
Scheduled daytime wheel-running activity in mice exposed to a skeleton photoperiod. (A) Experimental scheme: mice had free access to a wheel for 3 weeks under 12:12 L:D before transfer to a skeleton photoperiod (SPP). The SPP consists of 1h light pulse from ZT0 to ZT1 and from ZT11 to ZT12. After 3 weeks under SPP conditions, mice were separated into two groups: one with free access to wheel and food, the other with exclusive access to wheel during day and food during night. (B) Representative double-plotted actograms of DA mice, the gray background indicates light off, the arrow indicates the timeshift in wheel access. (C) 24-h activity levels of CTRL and DA mice, and (D) resulting average of day vs. night locomotor activity (D) Food intake during the active, feeding period. (E) 24-h core body temeprature levels of CTRL and DA mice, and (F) resulting day and night values. (G) Serum corticosterone levels. Data is shown as the average ± SEM (n = 24). * P < 0.05; ** P < 0.01; *** P < 0.001. Unpaired Student’s t-test (D, E, G) and One-way ANOVA (F).

Under SPP conditions, DA mice used the wheel from the first day of daytime-restricted access and maintained a stable onset and level of activity during the following days (**Fig. 5B, S4A**).

DA mice almost exclusively ran within the first 3h of daytime wheel access (**Fig. 5C**). Even though total activity only represented 50% of the control group (**Fig. 5D**), running intensity was comparable between the two groups during the first two hours of wheel access (**Fig. 5C, S4B**). Daytime-restricted wheel access promoted food intake in the first few hours of the nighttime period but significantly decreased the overall amount of food consumed (**Fig. 5E, S4C**). As expected, there was an elevation of daytime core body temperature paralleling the increase in wheel-running activity of DA mice (**Fig. 5F, G**). The temporal organization of nighttime core body temperature values in DA mice, however, closely resembled those of CTRL, nighttime active mice, likely caused by comparable feeding bout activities (**Fig. 5E**). To evaluate whether the animal handling, or the running at an abnormal time point caused a severe physiological stress, we measured serum corticosterone levels, but did not observe significant changes (**Fig. 5H**). This also demonstrates that daytime training does not affect the daily rhythm of corticosterone and, moreover, that voluntary wheel running is not associated with a stress response in mice, in stark contrast to treadmill exercise (**Fig. S1D**). This stress affects the clock and thereby confounds the analysis of Zeitgeber activity of exercise (Tahara et al., 2017).

### Daytime wheel running profoundly affects skeletal muscle gene expression but not the core clock

Having established a robust system of voluntary, stress-free daytime activity uncoupled from feeding behavior, we evaluated the impact of day- and nighttime activity on gene expression. To do so, we collected muscle tissues after 3 weeks of daytime wheel access, when running performance already plateaued in particular in DA mice (**Fig S4A**), without depriving mice of wheel access. First, we studied the impact of training on the daily expression of clock genes, which would reveal potential Zeitgeber activity. We have shown that acute bouts of treadmill exercise cause the rapid expression of some of the core clock transcriptional (co-)regulators (**Fig. 2C, Fig. S2C**). Moreover, nighttime wheel running exercise modulates the daily amplitude of core clock gene expression (Yasumoto et al., 2015). Prolonged daytime wheel running resulted in a marked effect on the amplitude, but not on the phase, of most core clock genes. For example, the relative amplitude (peak-trough amplitude divided by mean) of *Bmal1, Clock, Cry1-2, Per1-3, RORα* and *RORγ*, was dampened in muscles of the DA group (**Fig. 6A, S5A**).

**FIGURE 6.**
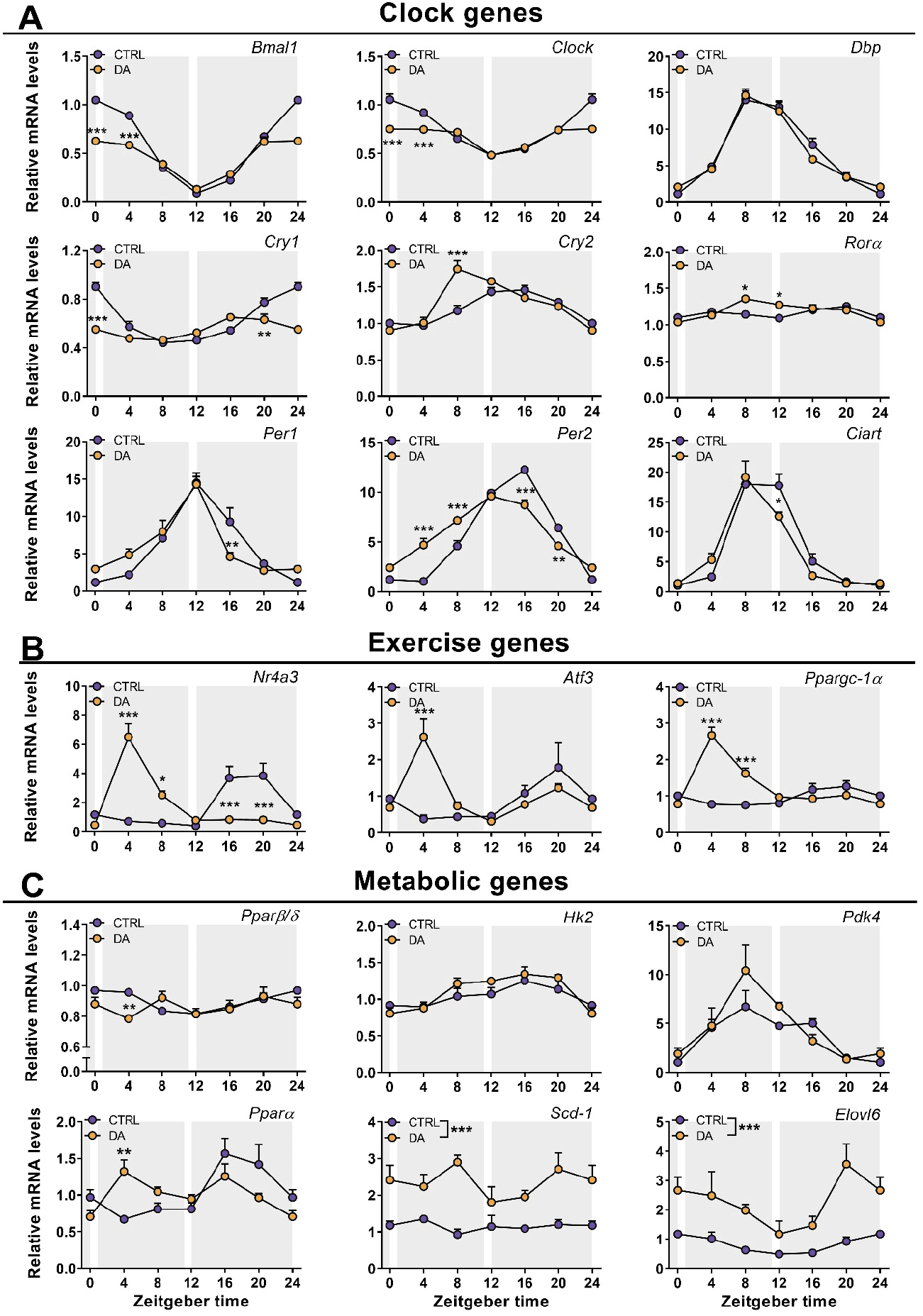
Daytime wheel running profoundly affects skeletal muscle gene expression but not the core clockwork. Gene expression in control (CTRL) and daytime activity (DA) mice. (A) Clock genes, (B) Metabolic genes, and (C) Exercise-related gene categories. Expression values were determined by qPCR and normalized to *Hprt*. Data is shown as the average fold-change ± SEM (n = 4) relative to the expression in CTRL ZT0 set to 1 * P < 0.05; ** P < 0.01; *** P < 0.001. Oneway ANOVA (A-C).

To elucidate whether daytime training affects the clock of other peripheral tissues, we also measured clock genes in the liver, without observing robust changes (**Fig. S6A**). Thus, to compare these effects to an established Zeitgeber, we finally determined clock expression in liver and muscle of mice undergoing daytime feeding (DF). As established (Damiola et al., 2000), this Zeitgeber paradigm resulted in a complete reversal of the liver clock (**Fig. S6B**). Interestingly, however, DF exerted an effect on the skeletal muscle clock that resembles that of daytime feeding, with a blunting of the amplitude of several genes, and minor phase shifts in others (Reznick et al., 2013) (**Fig. S6C**). Thus, collectively, the skeletal muscle clock seems much more refractory to external perturbations compared to the liver clock, even though training resulted in clear differences in the regulation of selected exercise (e.g., *Ppargc-1a*) and metabolic transcripts (e.g., *Ppara*) in muscles of DA mice (**Fig. 6B, C; S5B, C**).

### Characterization of the muscle transcriptome, proteome and phosphoproteome responses of daytime wheel active mice

We next dissected the global changes elicited by day-compared to nighttime activity on the transcriptome, proteome, and phosphoproteome at ZT4 and ZT16, near the peak of wheel-running activity in DA and CTRL mice, respectively. Daytime training led to the differential expression of 2094 genes at ZT4, with a similar proportion of down- and up-regulated transcripts (**Fig. 7A, S7**). In comparison, training at nighttime was a weaker modifier of gene expression: we only found 368 transcripts for which the expression was different between CTRL, nighttime-trained muscles and those obtained from daytime-trained animals at ZT16 (**Fig. 7A, S7**). The overlap between the day- and nighttime activity groups describes known exercise-regulated genes, including members of the KLF transcription factor gene family (i.e., *Klf4*, 5 and *15*), the MAF transcription factors and *Nr4a3*. Incidentally, *Atf3, Per2*, and *Ppargc-1a* were among the top induced genes only in daytime working muscles (**Fig.7B**). The nuclear receptor *Ppara*, a regulator of fatty acid oxidation (Muoio et al., 2002), was likewise specifically induced by daytime wheel running. Lastly, the KEGG pathways of DA mice reflected some of those that were activated by acute exercise bouts at early daytime, indicating a hardwiring of the pathways related to protein processing and mitophagy in chronic activity (**Fig. 7C, D**).

**FIGURE 7.**
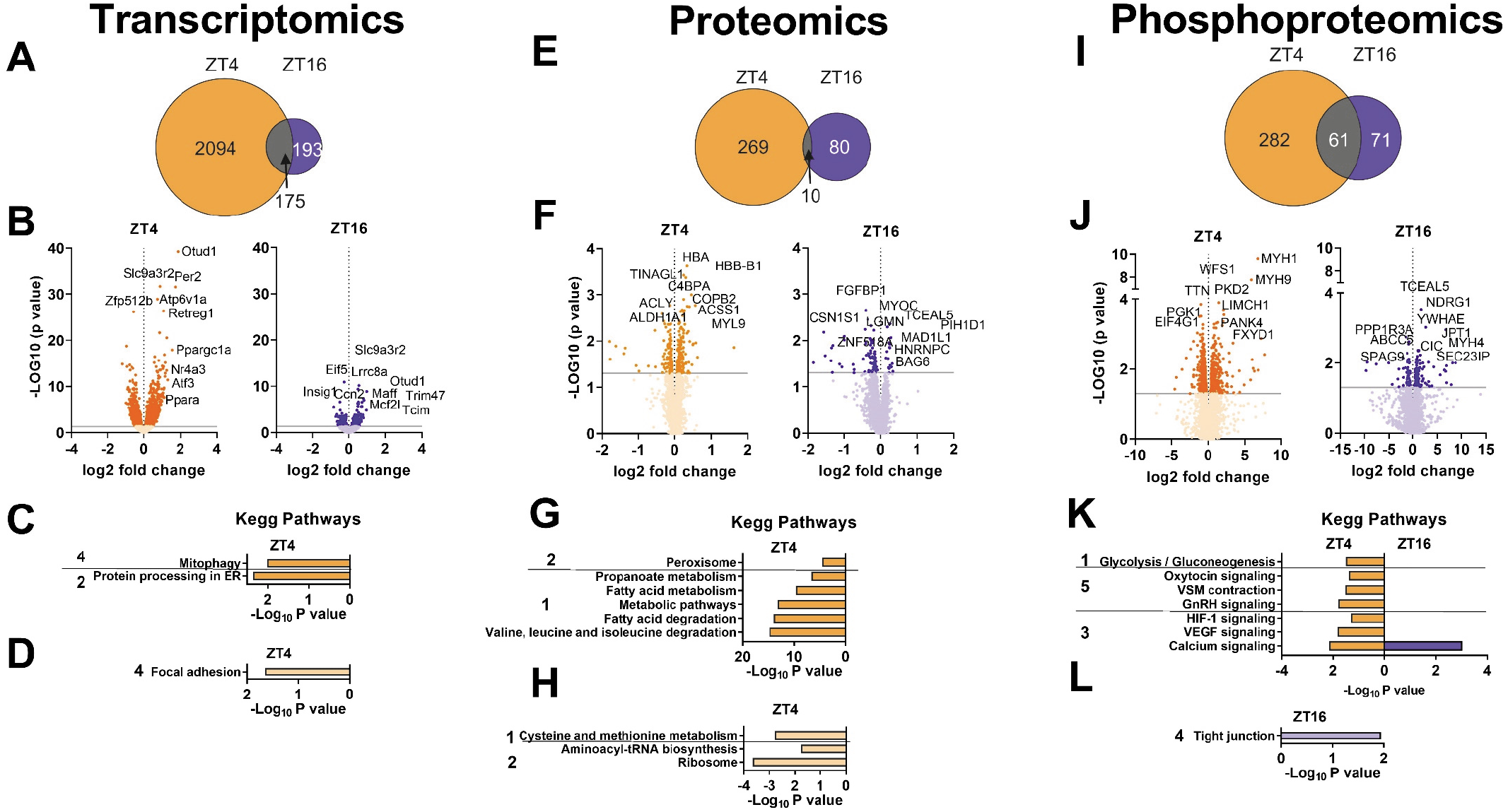
Characterization of the muscle transcriptome, proteome and phosphoproteome responses of daytime wheel active mice. (A) Venn diagrams with number of DEGs in DA mice at ZT4 (orange) and CTRL mice at ZT16 (blue), and resulting overlap (gray). (B) Volcano plots displaying the DEGs (top 10 shown) as identified in the same groups as above. KEGG analysis (top 5) of (C) up-regulated and (D) down-regulated genes as above. (E) Venn diagrams displaying the number of differentially regulated proteins for nighttime (blue), and daytime (orange) wheel running and the overlap (gray). (F) Volcano plot displaying the differentially regulated proteins (top 10 shown) by nighttime (blue) and daytime (orange) wheel running. KEGG analysis (top 5) for proteins (G) induced or (H) decreased by nighttime (blue) and daytime (orange) wheel running. (I) Venn diagram displaying the number of differentially phosphorylated proteins as described above. (J) Volcano plot displaying the differentially phosphorylated proteins (top 10 shown). KEGG analysis (top 5) for (K) phosphorylated and (L) dephosphorylated proteins. KEGG pathway categories: 1. Metabolism; 2. Genetic Information Processing; 3. Environmental Information Processing; 4. Cellular Processes; 5. Organismal Systems.

Up to 3800 proteins were detected in skeletal muscles of DA and CTRL mice, including 246 for which the expression was exclusively altered by daytime wheel running (**Fig. 7E, F**). The majority of up-regulated proteins were linked to metabolic functions (**Fig. 7G, H**), including an important cluster regulating fatty acid metabolism (e.g., ATP-citrate synthase (ACLY), mitochondrial trifunctional enzyme subunit (HADH)α, FASN and PDK4) and the citrate cycle (e.g., isocitrate dehydrogenase (IDH)2, succinate dehydrogenase (SDH)b) (**Fig. 7F**), most of which are direct targets of PPARα (Boergesen et al., 2012; Gan et al., 2018). In line with the transcriptomic data, exclusive nighttime running-induced proteins are more scarce, and not attributable to specific pathways by KEGG analysis. Moreover, in silico prediction of the secretome of both groups did not return notable myokine candidates. In regard to posttranslational modifications, we observed the differential phosphorylation of 344 and 135 proteins in response to daytime and nighttime training, respectively, with an overlap of 62 protein-specific posttranslational modifications, some of which associated with “calcium signaling” (**Fig. 7I**). However, most of the proteins shared between ZT4 and ZT16 were differentially phosphorylated by wheel running exercise. For example, CACNA1s was phosphorylated on Ser700 in the morning and dephosphorylated on Ser1617 in the evening. Similar results were found for RYR1 and STIM1, with the identification of previously uncharacterized sites. Notably, there was a strong daytime-dependent posttranslational regulation of PYGM on Ser15 (**Fig. 7J**), potentially promoting the degradation of glycogen (Johnson, 1992). Accordingly, “Glycolysis” was enriched in the DA group (**Fig. 7K, L**), and ALDOA, a key enzyme in the fourth step of glycolysis, was one of the top phosphorylated enzymes.

Overall, marked differences in gene expression, protein levels and phosphorylation patterns were observed in mice undergoing daytime compared to nighttime activity, implying a strong impact of the time of training on muscle physiology and function in absence of a major change in the regulation of the skeletal muscle clock.

## Discussion

Physical activity is a crucial behavior for which strong evolutionary pressure exists to ensure critical timing with availability of food and evasion of predators. Thus, a strong mutual interaction between skeletal muscle, the circadian timing system, and exercise has been suspected (Gutierrez-Monreal et al., 2020). First, the daily variations in metabolic and functional properties, ultimately controlled by the circadian clock, will affect endurance and strength capacities. Second, repeated aberrations from normal behavior, e.g., temporal changes in activity levels, could exert Zeitgeber activity to modulate and synchronize peripheral clocks according to altered environmental constraints. However, for both of these hypotheses, strong influence of time of day on muscle cellular responses and performance, and the Zeitgeber activity of exercise training, remain poorly described. Herein, we further evaluated these aspects in controlled animal studies, e.g., using SPPs as technical intervention to reduce confounding factors.

First, collectively, our results strongly suggest that training has very poor Zeitgeber activity on the skeletal muscle clock, barely affecting the phase of circadian gene expression in muscle, very different from the strong and paradigmatic effect of daytime feeding on the liver clock (Damiola et al., 2000) and **Fig. S6B**). In stark contrast, daytime training leads to a dramatically altered profile in terms of transcription, protein levels, and phosphoproteomics. In most aspects, the exercise response was amplified in the daytime activity group, and in part, resembled that following an acute exercise bout. For example, normal nighttime running wheel activity is linked to only a very moderate transcriptional response of *Ppargc-1α* gene expression. This gene responds very strongly to an acute exercise bout in naive mice, qualitatively and quantitatively comparable to what we have observed in the daytime running wheel group, which could be interpreted as a lack of training adaptation, or habituation, in this group of mice (Perry et al., 2010). Alternatively, it is conceivable that the difference in exercise response in daytime compared to nighttime running wheel activity is at least in part related to the difference in feeding state, since food access in both groups was restricted to the dark period. In particular, the difference in proteins related to fatty acid metabolism, citrate cycle, and glucose metabolism, indicates that the metabolic state in these two groups might instruct the response to a metabolic stimulus such as physical activity. It will be interesting to further dissect these variables by the addition of different time-restricted feeding interventions to the physical activity paradigm in future studies.

The running capacity of untrained mice at different times of the day, hence the influence of the clock on muscle function, will have to be interpreted in a similar manner. Metabolic constraints most likely contribute significantly to the observed differences in endurance observed in our present study as well as previously reported (Ezagouri et al., 2019). For example, we observed a strong correlation between performance and liver, and to a lower extent also skeletal muscle, glycogen. Depletion of glycogen stores is a major determinant of muscle fatigue in mice and humans, arguably liver glycogen even more in rodents than the skeletal muscle counterpart (Hargreaves and Spriet, 2018; Richter and Hargreaves, 2013). “Train low”, exercise under low glycogen conditions, is an established training paradigm in humans that, despite lower performance capacity in the individual endurance training bout, elicits a higher overall training adaptation (Hawley et al., 2018). Whether this is also the case in the mice running at ZT12 (lowest endurance performance, low liver and muscle glycogen stores) compared to those running at ZT0 (highest endurance performance, high liver and muscle glycogen stores) remains to be shown. Intriguingly, the transcriptomics, proteomics and phosphoproteomics analyses of these two different groups indicated that besides a robust, similar response of muscle to one acute exercise bout, very different pathways are activated depending on the time of the day. In the high performing mice running at ZT0, at pattern of immediate activation of pathways related to inflammation, the complement system and autophagy, followed by a later increase in ribosomes suggest an efficient repair and regeneration process. In parallel, a coordinated regulation of proteins involved in muscle contraction and energy provisioning, e.g., those involved in calcium signaling and GLUT4 vesicle transport, is observed. Finally, the increased expression of MUPs and asprosin could be interpreted as a muscle-derived signal to coordinate glycogenolysis and glucose release from liver. In stark contrast, the early response of skeletal muscle of the mice running at ZT12 indicates a broad activation of cellular stress, followed by an engagement of catabolic processes to match energetic demands in the absence of high abundance of storage of energy substrates, e.g., indicated by activation of AMPK and the glucocorticoid receptor, which could be interpreted as a signal for muscle protein degradation to fuel liver gluconeogenesis in this context. Accordingly, FoxO signaling is only observed in these mice. Differential control of proteostasis is furthermore underlined by the selective modulation of mTOR and related pathways.

The metabolic constraints that could explain the differences in performance depending on the time of the day are at least secondarily under the control of the intrinsic muscle clock, e.g., in regards to energy level. Future experiments with inducible, skeletal muscle-specific knockout models for clock components might thus help to clarify the contribution of this system to muscle metabolism and functionality if not confounded by other consequences of such dysregulations (Dyar et al., 2014). Our data demonstrate clear effects of acute exercise bouts, and of prolonged voluntary daytime activity on several components of the broader clock mechanism, indicating that at least some of the affected genes are regulated by exercise. It, however, is not clear how some of these proteins will impart such changes in level or activity on the clock, whether such perturbations are mitigated by other, unperturbed clock control mechanisms, and to what extent such proteins affect other cellular processes besides clock oscillation. For example, the *Period 1-2* genes, which we found to be robustly induced by exercise at different times of the day, have been described as integrators of external and internal perturbations at the intersection of modulating the molecular clock and other cellular functions (Ripperger and Albrecht, 2012). Based on our data, these systems can become unlinked in certain settings and contexts, even though mutual regulation has been found, e.g., also by AMPK, HIF1α, or mTOR on the clock (Panda, 2016; Reinke and Asher, 2019).

Our data also indicate that appropriate timing of exercise bouts might facilitate repair and regeneration, important to speed up recovery and potentially allow higher intensity and/or volume. The best time of training, however, will not be deducible from experiments performed in nocturnal, in-breed laboratory mouse strains that do not recapitulate the complexity of the situation in humans with genetic variations, chronotypes, difference in eating habits and sleeping patterns, and other factors. Thus, athletes will have to be assessed individually (“personalized training”). Our comprehensive transcriptome, proteome, and phosphoproteome analyses might provide a starting point to identify and validate markers for such a stratification in humans. Lastly, our data indicate that training is not only not a strong Zeitgeber in a healthy mouse muscle, but, if chronically performed at the wrong time of the day, might lead to perturbations of the muscle clock which resemble perturbations of other peripheral clocks in pathological states and in aging, characterized by a general dampening of oscillation amplitudes (Chellappa et al., 2019; Rijo-Ferreira and Takahashi, 2019). Our experiments, however, will not allow any conclusions about a potential therapeutic effect of timed training (chrono-exercise) on muscle and other peripheral clocks in pathologies that are linked to abnormal circadian rhythmicity (Cederroth et al., 2019). Regardless of the outcome of future studies in such a direction, however, the negative results of our studies in regard to the effect of daytime activity in mice as Zeitgeber of the muscle clock should not be interpreted to discount training interventions, which have an established efficacy in preventing and treating a number of chronic diseases, often even rivaling or even surpassing that of currently available drugs (Pedersen and Saltin, 2015).

### Limitations of the study

We have used two different exercise modalities, in acute and chronic modes, and thus did not directly compare our multi-omics datasets. Multisite phosphorylation is a widespread phenomenon in eukaryotic cells, and a vast number of phosphorylated sites identified in our study are not yet associated with specific functional characteristics (e.g., enzymatic activity, binding, cellular localization or stability (Salazar and Hofer, 2009; Suwanmajo and Krishnan, 2015). We therefore have mainly limited our comparative phosphorylome analyses to proteins showing time- and exercise-dependent phosphorylation changes on unique residues. We, however, hope that the different responses of individual sites to timed exercise will provide valuable resources for further experimental investigation.

## Materials and methods

### Animals

8 weeks old C57BL/6JRj male mice (Janvier Labs) were housed in standard cages under 12:12 light:dark (LD) conditions, with light onset at 6am (Zeitgeber Time 0; ZT0) or entrained to a skeleton photoperiod (SPP) as described below and in Fig. 4, unless otherwise stated. The mice had ad libitum access to a standard chow diet (Maintenance 3432, KLIBA NAFAG) and water, unless otherwise stated. All experiments were performed in agreement with the principles of the Basel Declaration, with Federal and Cantonal Laws regulating the care and use of experimental animals in Switzerland, and the institutional guidelines of the Biozentrum and the University of Basel. The protocol with all methods described here was approved by the “Kantonales Veterinäramt” of the Kanton Basel-Stadt, under consideration of the well-being of the animals and the 3R principle.

### Forced high-intensity exercise performance across the day

Sedentary mice were acclimated to the treadmill equipped with a shock grid (Columbus Instruments, Columbus, Ohio, USA) on three consecutive days prior to the experiment. The accommodation period consisted in: Day 1, placing the mice in the treadmill for 10 min without shock and belt movement followed by 5 min at 5 m/min; Day 2, running at 5 m/min, 7 m/min and 10 m/min for 5 min each, without shock; Day 3, running at 8 m/min, 10 m/min and 12 m/min for 5 min each, with shock. After one resting day, a maximal exercise capacity test was performed by 3min at 8m/min increasing treadmill speed by 2 m every 2 min, at a 15° slope, until exhaustion. To provide additional motivation to avoid the lower portion of the treadmill and thus the shock grid, we gently scratched the back of the animal during accommodation and maximal exercise capacity test. Exhaustion was met if an animal remained on the electrical grid (providing a mild electrical stimulus of 0.5 mA, 200 ms pulse, 1 Hz) for more than 5 s. Tail blood glucose (Accu-Chek, Roche) and lactate (Lactate Plus meter, Nova Biomedical) values were determined immediately prior to treadmill exercise and within 1 min after physical exhaustion. Mice were either sacrificed immediately after exhaustion (≤ 5 min of time delay; Ex+0h) or 3h after exercise (Ex+3h). For the latter group, mice were returned to their home cages without access to food. This experiment was repeated every 4h for 24h starting from ZT0 (6 a.m.; scheme in Fig. 1 (A)). A non-exercised group (Sedentary; Sed) of mice was always sacrificed at a similar ZT (≤ 30 min of time delay) as the exercised mice (Ex+0h and Ex+3h). Note that control Sed mice were placed in new cages, with new bedding but no food access 60 min prior to sacrifice. Importantly, to take into account changes in basal gene expression over time, gene expression data obtained from Ex+0h and Ex+3h mouse groups running, e.g., at ZT0, were compared to sedentary controls sacrificed at ZT0 and ZT4, respectively

### Daytime scheduled wheel-running activity

Mice were single-housed in standard cages, within an environment-controlled cabinet (UniProtect Air Flow Cabinet, Bioscape), with the temperature set to 23°C. The mice had access to wheel with rods (ø 11.5 cm, Starr Life Sciences) under constant 12:12 LD conditions for three weeks prior to exposure to a skeleton photoperiod (SPP). The SPP consisted of two repeated light-pulses (LP): 1 h LP at the beginning of the resting period and 1 h LP at the end of the resting period, interrupted by 10h of darkness (Fig. 4A). After three weeks acclimatization to the SPP, three weeks of intervention followed. During the intervention, one group of mice (control, CTRL) had ad libitum access to food, water and free access to running wheels. Another group of mice (Daytime activity, DA) was food-restricted to the longer dark phase (active phase) and had access to a wheel only during the shorter dark period (resting phase) (see Fig. 4). Wheel and food access were controlled manually, without opening the cage to not disturb the animals. Importantly, the use of the wheel (light phase: 309 turns vs. dark phase: 17732 turns) and food intake (light phase: 0.4g vs. dark phase: 4.7g) during the resting/inactive period was virtually absent in the CTRL group. Lastly, we only compared daytime and nighttime wheel active animals in our study, trained for a similar length of time; the comparison of sedentary mice with mice given free access to a running wheel is well documented elsewhere (Allen et al., 2001; Holloszy, 1967; McKie et al., 2019).

### Body temperature and locomotor activity recordings

General locomotor activity and core body temperature data were acquired with the E-Mitter Telemetry System (Starr Life Sciences) from single-caged animals placed in an environment-controlled cabinet (UniProtect Air Flow Cabinet, Bioscape). Briefly, small transponders (G2 EMitter, Starr Life Sciences) were implanted into the abdominal cavity of mice under isoflurane anesthesia (2 % isoflurane + O2). Mice were treated with Meloxicam (1 mg/kg) pre- and post-operatively and allowed to recover for three weeks. The abovementioned parameters together with the wheel-running activity were recorded with a PC-based acquisition system connected to ER4000 Receivers (VitalView, Starr Life Sciences).

### Muscle tissue preparation and blood collection

Mice from the different experiments were sacrificed by short exposure to CO2 and immediate exsanguination. Blood was collected in tubes containing lithium heparin (Microvette 500 LH, Sarstedt, 20.1345) centrifuged at 2000 g for 5 min at RT and stored at −80°C. The glycolytic quadriceps and gastrocnemius muscles, as well as liver samples were quickly snap frozen in liquid nitrogen and stored at −80°C until further analysis.

### Quantitative Real-Time PCR (Reverse Transcript)

Total RNA from muscle tissues was extracted using a hybrid method combining TRI-Reagent (Sigma-Aldrich T9424) and RNeasy Mini Kit (QIAGEN 74104). RNA quantity and purity were measured with a NanoDrop OneC (ThermoFisher Scientific). High-Capacity cDNA Reverse Transcript Kit (Applied Biosystems, 4368814) was used for cDNA synthesis with 1 ug of total RNA. Quantitative real-time PCR was performed with Fast SYBR Green Master Mix (Applied Biosystems, 4385612) in a RT-PCR System (StepOnePlus, Applied Biosystems). PCR reactions were done in duplicate with the addition of negative controls (i.e., no reverse transcription and no template controls). Relative expression levels were determined using the comparative ΔΔCT method to normalize target gene mRNA to *Hprt*. Rhythmicity and differential rhythmicity were assessed using the methods RAIN (Thaben and Westermark, 2014) and DODR (Thaben and Westermark, 2016), respectively.

### Blood parameters analysis

Quantification of plasma triglyceride was done with the Cobas c111 analyzer (Roche). Plasma free fatty acids were analyzed using the Free Fatty Acid Quantification Assay Kit (Abcam, ab65341) following manufacturer’s recommendations. Muscle and liver glycogen levels were measured with the Glycogen Assay Kit (Abcam, ab65620).

### RNA sequencing and data analysis

RNA quality was determined on the Bioanalyzer instrument (Agilent Technologies, Santa Clara, CA, USA) using the RNA 6000 Nano Chip (Agilent, Cat# 5067-1511) and quantified by Spectrophotometry using the NanoDrop ND-1000 Instrument (NanoDrop Technologies, Wilmington, DE, USA). Library preparation was performed with 1μg total RNA using the TruSeq Stranded mRNA Library Prep Kit High Throughput (Cat# RS-122-2103, Illumina, San Diego, CA, USA). Libraries were quality-checked on the Fragment Analyzer (Advanced Analytical, Ames, IA, USA) using the Standard Sensitivity NGS Fragment Analysis Kit (Cat# DNF-473, Advanced Analytical) revealing excellent quality of libraries (average concentration was 152±9 nmol/L and average library size was 374±4 base pairs). Samples were pooled to equal molarity. Each pool was quantified by PicoGreen Fluorometric measurement in order to be adjusted to 1.8pM and used for clustering on the NextSeq 500 instrument (Illumina). Samples were sequenced Single-reads 76 bases using the NextSeq 500 High Output Kit 75-cycles (Illumina, Cat# FC-404-1005), and primary data analysis was performed with the Illumina RTA version 2.4.11 and Basecalling Version bcl2fastq-2.20.0.422.

To quantify mRNA expression levels, kallisto version 0.46.0 (Bray et al., 2016) was used. To build the index for kallisto, the GRCm38.p6 (mm10) genome assembly and the ncbiRefSeqCurated transcript annotation of the UCSC genome browser were used (Karolchik et al., 2014; Pruitt et al., 2014). microRNAs (miRbase Version 19 (Kozomara and Griffiths-Jones, 2011), translated to RefSeq IDs through BioMart (Durinck et al., 2009)) were excluded. Only one transcript was retained if several had both identical start and end coordinates, slightly flattening the annotation, preference was given to transcripts with IDs starting with “NM_”. Transcripts mapping to more than one chromosome, or to random or chrUn contigs, were discarded. tximport version 1.14.0 (Soneson et al., 2015) was used to transform expression levels to flattened gene level pseudo-counts, using the “lengthScaledTPM” option. For this, RefSeq IDs were mapped to Entrez gene IDs using the org.Mm.eg.db database of R/Bioconductor version 3.10 (Gentleman et al., 2004). DESeq2 version 1.26.0 (Love et al., 2014) was used for statistical analysis of gene level differential expression. Here, log2 fold changes were estimated by the DESeq2 shrinkage estimator.

### Proteomics and data analysis

Phosphopeptide enrichment and LC MS/MS analysis

Tissue was lysed in 8M Urea, 0.1M ammonium bicarbonate, phosphatase inhibitors (Sigma P5726&P0044) by sonication (Bioruptor, 10 cycles, 30 seconds on/off, Diagenode, Belgium) and proteins were digested as described previously (Ahrne et al., 2016). Shortly, proteins were reduced with 5 mM TCEP for 60 min at 37 °C and alkylated with 10 mM chloroacetamide for 30 min at 37 °C. After diluting samples with 100 mM ammonium bicarbonate buffer to a final urea concentration of 1.6M, proteins were digested by incubation with sequencing-grade modified trypsin (1/50, w/w; Promega, Madison, Wisconsin) for 12 h at 37°C. After acidification using 5% TFA, peptides were desalted using C18 reverse-phase spin columns (Macrospin, Harvard Apparatus) according to the manufacturer’s instructions, dried under vacuum and stored at −20°C until further use.

Peptide samples were enriched for phosphorylated peptides using Fe(III)-IMAC cartridges on an AssayMAP Bravo platform as recently described (Post et al., 2017). Unmodified peptides (“flowthrough”) were subsequently used for TMT analysis.

Phospho-enriched peptides were resuspended in 0.1% aqueous formic acid and subjected to LC–MS/MS analysis using a Q Exactive HF Mass Spectrometer or an Orbitrap Fusion Lumos Mass Spectrometer fitted with an EASY-nLC 1000 or an EASY-nLC 1200, respectively (both Thermo Fisher Scientific) and a custom-made column heater set to 60°C. Peptides were resolved using a RP-HPLC column (75μm × 30cm or 75μm × 36cm, respectively) packed inhouse with C18 resin (ReproSil-Pur C18–AQ, 1.9 μm resin; Dr. Maisch GmbH) at a flow rate of 0.2 μl/min. The following gradient was used for peptide separation: Q Exactive HF from 5% B to 8% B over 5 min to 20% B over 45 min to 25% B over 15 min to 30% B over 10 min to 35% B over 7 min to 42% B over 5 min to 50% B over 3min to 95% B over 2 min followed by 18 min at 95% B, Orbitrap Fusion Lumos from 5% B to 8% B over 5 min to 20% B over 45 min to 25% B over 15 min to 30% B over 10 min to 35% B over 7 min to 42% B over 5 min to 50% B over 3min to 95% B over 2 min followed by 18 min at 95% B. Buffer A was 0.1% formic acid in water and buffer B was 80% acetonitrile, 0.1% formic acid in water.

The Q Exactive HF mass spectrometer was operated in DDA mode with a total cycle time of approximately 1 s. Each MS1 scan was followed by high-collision-dissociation (HCD) of the 10 most abundant precursor ions with dynamic exclusion set to 45 seconds. For MS1, 3e6 ions were accumulated in the Orbitrap over a maximum time of 100 ms and scanned at a resolution of 120,000 FWHM (at 200 m/z). MS2 scans were acquired at a target setting of 1e5 ions, maximum accumulation time of 100 ms and a resolution of 30,000 FWHM (at 200 m/z). Singly charged ions and ions with unassigned charge state were excluded from triggering MS2 events. The normalized collision energy was set to 28%, the mass isolation window was set to 1.4 m/z and one microscan was acquired for each spectrum. The Orbitrap Fusion Lumos mass spectrometer was operated in DDA mode with a cycle time of 3 seconds between master scans. Each master scan was acquired in the Orbitrap at a resolution of 120,000 FWHM (at 200 m/z) and a scan range from 375 to 1600 m/z followed by MS2 scans of the most intense precursors in the Orbitrap at a resolution of 30,000 FWHM (at 200 m/z) with isolation width of the quadrupole set to 1.4 m/z. Maximum ion injection time was set to 50ms (MS1) and 54 ms (MS2) with an AGC target set to 1e6 and 5e4, respectively. Only peptides with charge state 2 – 5 were included in the analysis. Monoisotopic precursor selection (MIPS) was set to Peptide, and the Intensity Threshold was set to 2.5e4. Peptides were fragmented by HCD (Higher-energy collisional dissociation) with collision energy set to 30%, and one microscan was acquired for each spectrum. The dynamic exclusion duration was set to 30s.

The acquired raw-files were imported into the Progenesis QI software (v2.0, Nonlinear Dynamics Limited), which was used to extract peptide precursor ion intensities across all samples applying the default parameters. The generated mgf-file was searched using MASCOT against a murine database (consisting of 34026 forward and reverse protein sequences downloaded from Uniprot on 20190129) and 392 commonly observed contaminants using the following search criteria: full tryptic specificity was required (cleavage after lysine or arginine residues, unless followed by proline); 3 missed cleavages were allowed; carbamidomethylation (C) was set as fixed modification; oxidation (M) and phosphorylation (STY) were applied as variable modifications; mass tolerance of 10 ppm (precursor) and 0.02 Da (fragments). The database search results were filtered using the ion score to set the false discovery rate (FDR) to 1% on the peptide and protein level, respectively, based on the number of reverse protein sequence hits in the datasets. Exported peptide intensities were normalized based on the protein regulations observed in the corresponding TMT experiments in order to account for changes in protein abundance. Only peptides corresponding to proteins, which were regulated significantly with a p value ≤ 5% in the TMT analysis were normalized. Quantitative analysis results from label-free quantification were processed using the SafeQuant R package v.2.3.2. (Ahrne et al., 2016), https://github.com/eahrne/SafeQuant/) to obtain peptide relative abundances. This analysis included global data normalization by equalizing the total peak/reporter areas across all LC-MS runs, data imputation using the knn algorithm, summation of peak areas per and LC–MS/MS run, followed by calculation of peptide abundance ratios. Only isoform specific peptide ion signals were considered for quantification. The summarized peptide expression values were used for statistical testing of between condition differentially abundant peptides. Here, empirical Bayes moderated t-Tests were applied, as implemented in the R/Bioconductor limma package (http://bioconductor.org/packages/release/bioc/html/limma.html) were used.TMT labelling and LC MS/MS analysis

Tryptic peptides were labeled with isobaric tandem mass tags (TMT10plex or TMTpro 16plex, Thermo Fisher Scientific). Peptides were resuspended in labeling buffer (2 M urea, 0.2 M HEPES, pH 8.3) by sonication and TMT reagents were added to the individual peptide samples followed by a 1 h incubation at 25°C shaking at 500 rpm. To quench the labelling reaction, aqueous 1.5 M hydroxylamine solution was added and samples were incubated for another 5 min at 25°C shaking at 500 rpm followed by pooling of all samples. The pH of the sample pool was increased to 11.9 by adding 1 M phosphate buffer (pH 12) and incubated for 20 min at 25°C shaking at 500 rpm to remove TMT labels linked to peptide hydroxyl groups. Subsequently, the reaction was stopped by adding 2 M hydrochloric acid until a pH < 2 was reached. Finally, peptide samples were further acidified using 5 % TFA, desalted using Sep-Pak Vac 1cc (50 mg) C18 cartridges (Waters) according to the manufacturer’s instructions and dried under vacuum. For TMTpro 16plex analysis, 4 peptide samples were prepared from C2C12 cells, TMT labelled and included in the analysis to boost protein coverage.

TMT-labeled peptides were fractionated by high-pH reversed phase separation using a XBridge Peptide BEH C18 column (3,5 μm, 130 Å, 1 mm x 150 mm, Waters) on an Agilent 1260 Infinity HPLC system. Peptides were loaded on column in buffer A (20 mM ammonium formate in water, pH 10) and eluted using a two-step linear gradient from 2% to 10% in 5 minutes and then to 50% buffer B (20 mM ammonium formate in 90% acetonitrile, pH 10) over 55 minutes at a flow rate of 42 μl/min. Elution of peptides was monitored with a UV detector (215 nm, 254 nm) and a total of 36 fractions were collected, pooled into 12 fractions using a post-concatenation strategy as previously described (Wang et al., 2011) and dried under vacuum.

Dried peptides were resuspended in 0.1% aqueous formic acid and subjected to LC–MS/MS analysis using a Q Exactive HF Mass Spectrometer or an Orbitrap Fusion Lumos Mass Spectrometer fitted with an EASY-nLC 1000 or an EASY-nLC 1200, respectively (both Thermo Fisher Scientific) and a custom-made column heater set to 60°C. Peptides were resolved using a RP-HPLC column (75μm × 30cm or 75μm × 36cm, respectively) packed inhouse with C18 resin (ReproSil-Pur C18–AQ, 1.9 μm resin; Dr. Maisch GmbH) at a flow rate of 0.2 μLmin-1. The following gradient was used for peptide separation: Q Exactive HF from 5% B to 15% B over 10 min to 30% B over 60 min to 45 % B over 20 min to 95% B over 2 min followed by 18 min at 95% B, Orbitrap Fusion Lumos from 5% B to 15% B over 9 min to 30% B over 90 min to 45 % B over 21 min to 95% B over 2 min followed by 18 min at 95% B. Buffer A was 0.1% formic acid in water and buffer B was 80% acetonitrile, 0.1% formic acid in water.

The Q Exactive HF mass spectrometer was operated in DDA mode with a total cycle time of approximately 1 s. Each MS1 scan was followed by high-collision-dissociation (HCD) of the 10 most abundant precursor ions with dynamic exclusion set to 30 seconds. For MS1, 3e6 ions were accumulated in the Orbitrap over a maximum time of 100 ms and scanned at a resolution of 120,000 FWHM (at 200 m/z). MS2 scans were acquired at a target setting of 1e5 ions, maximum accumulation time of 100 ms and a resolution of 30,000 FWHM (at 200 m/z). Singly charged ions and ions with unassigned charge state were excluded from triggering MS2 events. The normalized collision energy was set to 30%, the mass isolation window was set to 1.1 m/z and one microscan was acquired for each spectrum. The Orbitrap Fusion Lumos mass spectrometer was operated in DDA mode with a cycle time of 3 seconds between master scans. Each master scan was acquired in the Orbitrap at a resolution of 120,000 FWHM (at 200 m/z) and a scan range from 375 to 1600 m/z followed by MS2 scans of the most intense precursors in the Orbitrap at a resolution of 30,000 FWHM (at 200 m/z) with isolation width of the quadrupole set to 1.1 m/z. Maximum ion injection time was set to 50ms (MS1) and 54 ms (MS2) with an AGC target set to 1e6 and 1e5, respectively. Monoisotopic precursor selection (MIPS) was set to Peptide, and the Intensity Threshold was set to 5e4. Peptides were fragmented by HCD (Higher-energy collisional dissociation) with collision energy set to 38%, and one microscan was acquired for each spectrum. The dynamic exclusion duration was set to 45s.

The acquired raw-files were analysed using the SpectroMine software (Biognosis AG, Schlieren, Switzerland). Spectra were searched against a murine database consisting of 17013 protein sequences (downloaded from Uniprot on 20190307) and 392 commonly observed contaminants. Standard Pulsar search settings for TMT10plex (“TMT10plex Quantification”) and TMTpro 16plex (“TMTpro Quantification”) were used and resulting identifications and corresponding quantitative values were exported on the PSM level using the “Export Report” function. Acquired reporter ion intensities in the experiments were employed for automated quantification and statistical analysis using our in-house developed SafeQuant R script v2.3.2, (Ahrne et al., 2016). This analysis included adjustment of reporter ion intensities, global data normalization by equalizing the total reporter ion intensity across all channels, summation of reporter ion intensities per protein and channel, calculation of protein abundance ratios and testing for differential abundance using empirical Bayes moderated t-statistics.

### Pathway enrichment analysis

For the pathway enrichment analysis with g:Profiler (Reimand et al., 2016), all genes and (phospho-)proteins of a time point were used, including the overlap.

### Statistics

The n number used per genotype for each experiment is indicated in the figure legend. Data are represented as mean±SEM and statistically analyzed with GraphPad Prism 8. Two-tailed, nonpaired Student t-test was performed to evaluate statistical difference between two groups. For multiple comparisons, data were analyzed using 2-Way ANOVA followed by Sidak’s multiple comparisons test. Corresponding symbols to highlight statistical significance are the following: *: p≤0.05; **: p≤0.01; ***: p≤0.001.

## Acknowledgments

We are very grateful to the Quantitative Genomics Facility, in particular Phillippe Demougin for preparing cDNA libraries. We thank Katja Lamia, Scripps Research, for helpful comments on our manuscript. We also acknowledge the support of the Center for Scientific Computing (sciCORE) of the University of Basel.

## Author contributions

Conceptualization and supervision, J.D. and C.H.; Methodology, G.M., J.D., P.O.W., D.R; Investigation, G.M., J.D., G.S.; Formal analysis and visualization, G.M., J.D., P.O.W., D.R; Writing – Original Draft, G.M, J.D.; Writing – Review & Editing, G.M, J.D., P.O.W., C.H.; Funding Acquisition, C.H.

## Data deposition

The data have been deposited in GEO (accession numbers XXX) and.....

## Abbreviations

AMPK: AMP-activated protein kinase
Atf3: activating transcription factor 3
Bmal1: brain and muscle ARNT (aryl hydrocarbon receptor nuclear translocator)-Like 1
Ciart: circadian associated repressor of transcription
Clock: circadian locomotor output cycles kaput
Cry 1/2/3: crypochrome1/2
Ctrl: control
DA: daytime activity
Dbp: albumin D box-binding protein
GLUT4: glucose transporter 4
Hk2: hexokinase 2
LD: light-dark
Nr4a3: nuclear receptor subfamily 4 group A member 3 / NOR-1
Pdk4: pyruvate dehydrogenase kinase 4
Per1/2/3: period 1/2/3
Ppargc-1α / PGC-1α: peroxisome proliferator-activated receptor γ coactivator 1α
SPP: skeleton photoperiod
Tbc1d1: TBC1 domain family member 1
ZT: Zeitgeber time

