## Supplemental Figures 1-7 for "Transcriptomic, proteomic and phosphoproteomic underpinnings of daily exercise performance and Zeitgeber activity of endurance training"

**Supplemental Information**

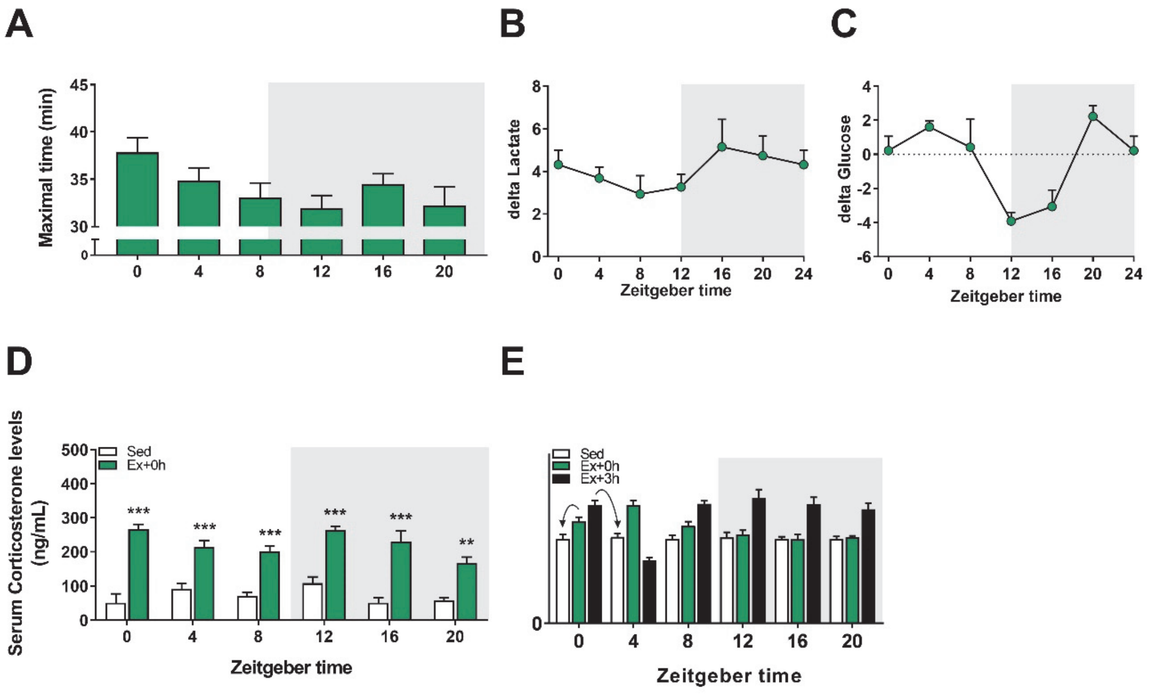


**SUPPL. FIGURE 1. Time-of-day-dependent variations in mouse treadmill exercise performance.**

(A) Maximal time reached at exhaustion. (B) Delta blood lactate and (C) delta glucose levels (resting values subtracted from values at exhaustion). (D) Serum corticosterone levels in Sed and at exhaustion. Data is shown as the average ± SEM (n=3 per group and time point). * P < 0.05; ** P < 0.01; *** P < 0.001. One-way ANOVA (D). (E) Example for the analysis of the bar graph plots, arrows indicate normalization. Note, values are plotted according to their exercise time, while the normalization was performed corresponding to their sacrifice time (i.e., the gene expression of a mouse trained at ZT0 and sacrificed directly at exhaustion was normalized to the expression of Sed ZT0 plotted at ZT0, while the gene expression of a mouse trained at ZT0 and sacrificed 3h after exhaustion was normalized to the expression of Sed ZT4, but plotted at ZT0, at its training time).

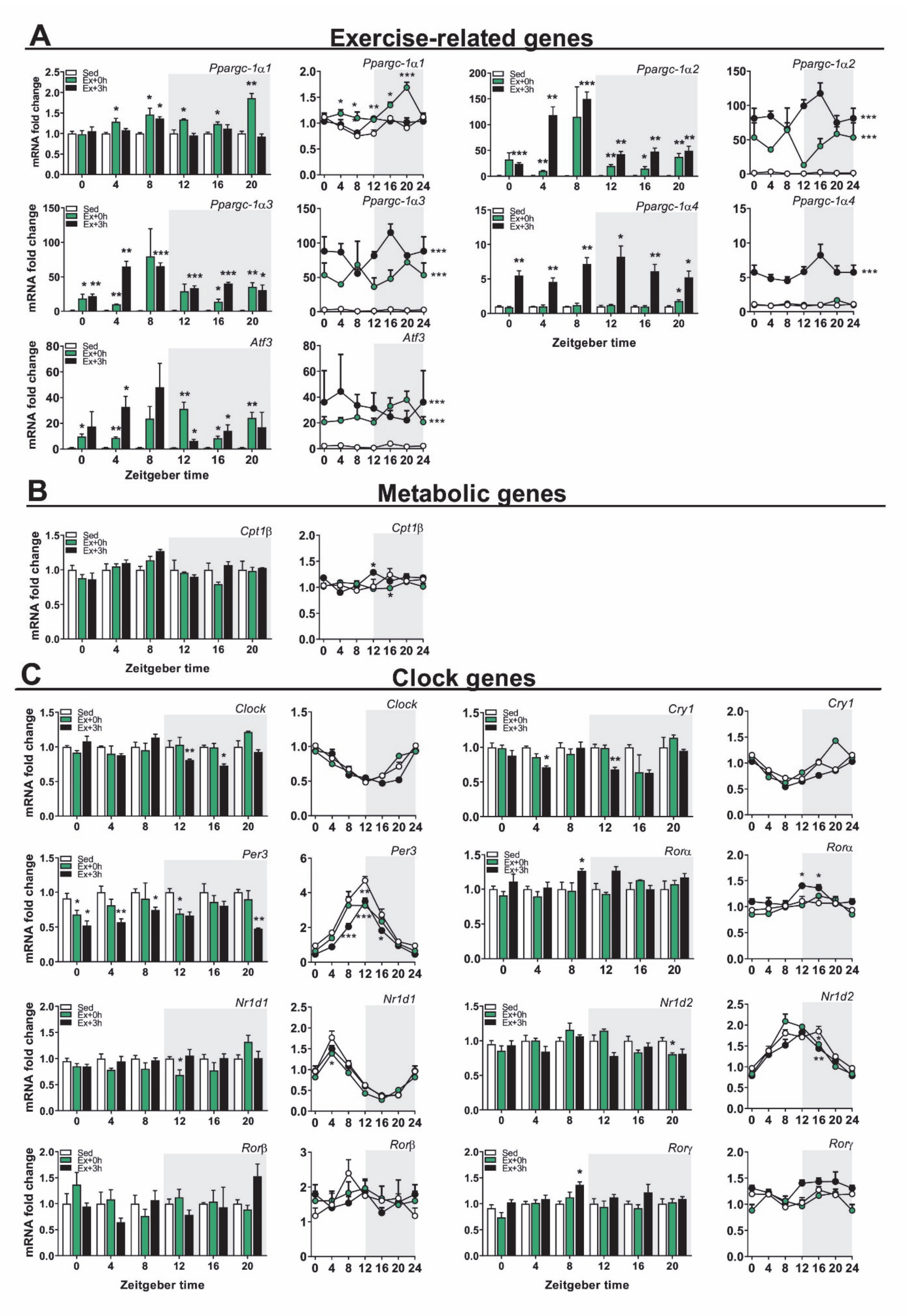


**SUPPL. FIGURE 2. Scheduled treadmill induces broad and time-dependent transcriptional responses in skeletal muscle.**

Gene expression in sedentary mice (Sed) and exercised mice at (Ex+0h) and 3h (Ex+3h) after exhaustion. (A) Exercise-related genes, (B) Metabolic genes, (C) Clock genes categories. Expression values were determined by qPCR and normalized to *Hprt*. Data in bar graph is shown as the average fold-change ± SEM (n = 3) relative to the expression in Sed set to 1 (see method for details on normalization). Data in line graph is shown as the average fold-change ± SEM (n = 3) relative to the expression in the Sed ZT0 group set to 1. * $P < 0.05$; ** $P < 0.01$; *** $P < 0.001$. Unpaired Student's t-test (Bar graphs) and One-way ANOVA (Line graphs).

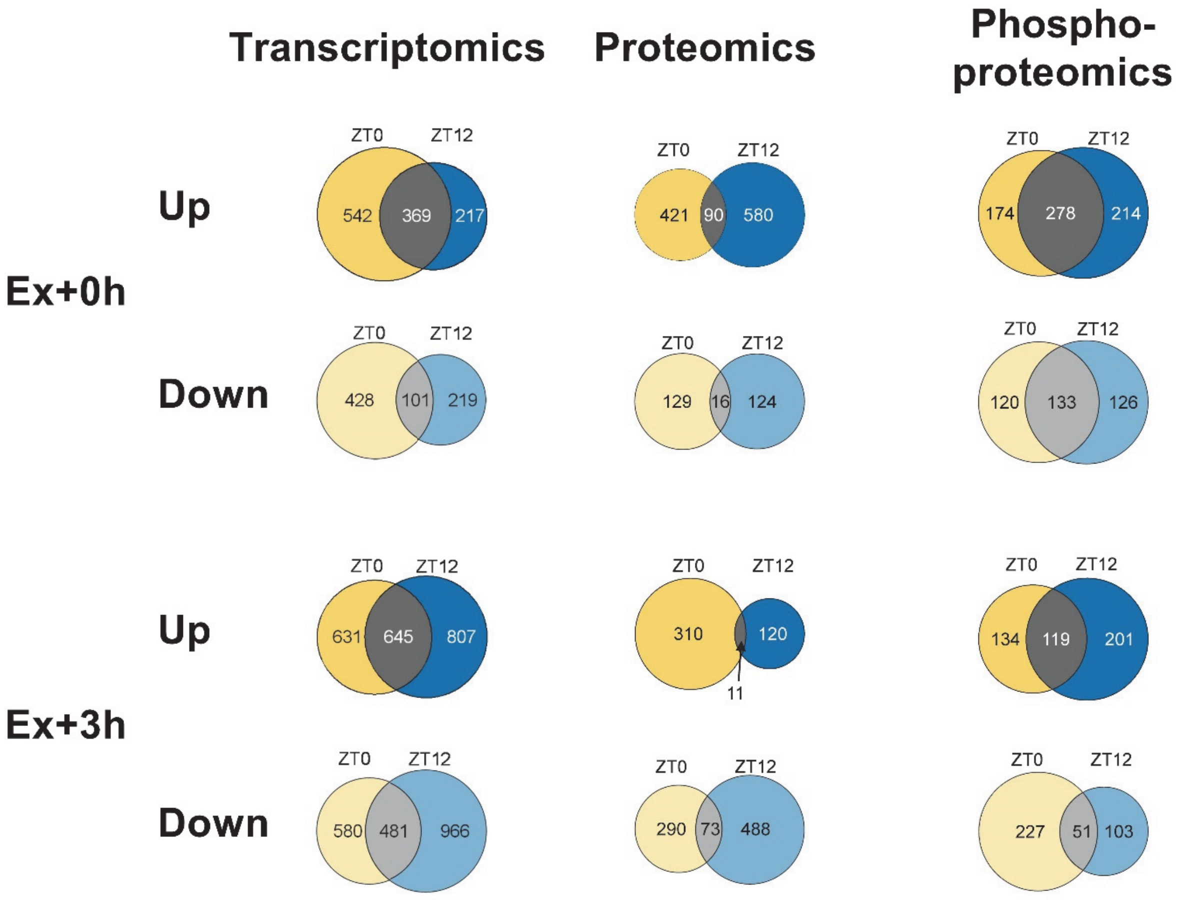


**SUPPL. FIGURE 3. Distinct muscle gene expression signatures after early light vs. early dark phase treadmill exercise.**

Venn diagrams displaying the number of differentially regulated genes, proteins and phosphosites immediately (A; Ex+0h) or 3h (B; Ex+3h) after early daytime (ZT0; yellow), early nighttime (ZT12; blue) exercise, and the overlap (gray).

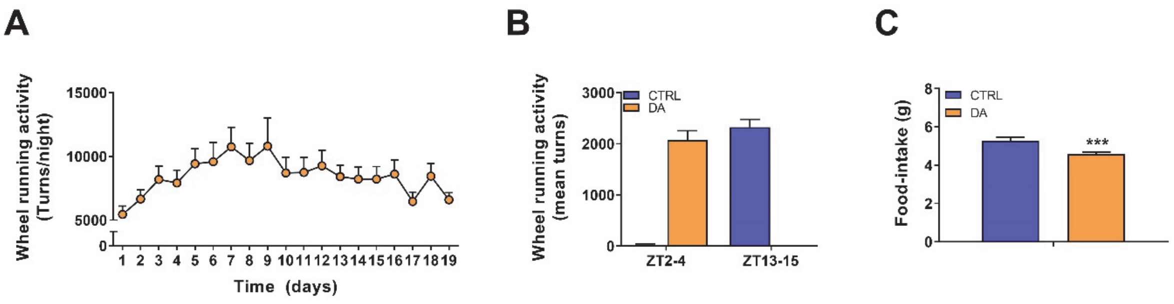


**SUPPL. FIGURE 4. Physiological parameters of daytime active mice.**

(A) Daily wheel-running activity from the first day of restriction of the DA mice. Data is shown as the average fold-change ± SEM (n = 24). (B) Quantified wheel-running activity during the first two hours of wheel excess. (C) Food-intake average over the full food access period. *** P < 0.001. Unpaired Student's t-test (C).

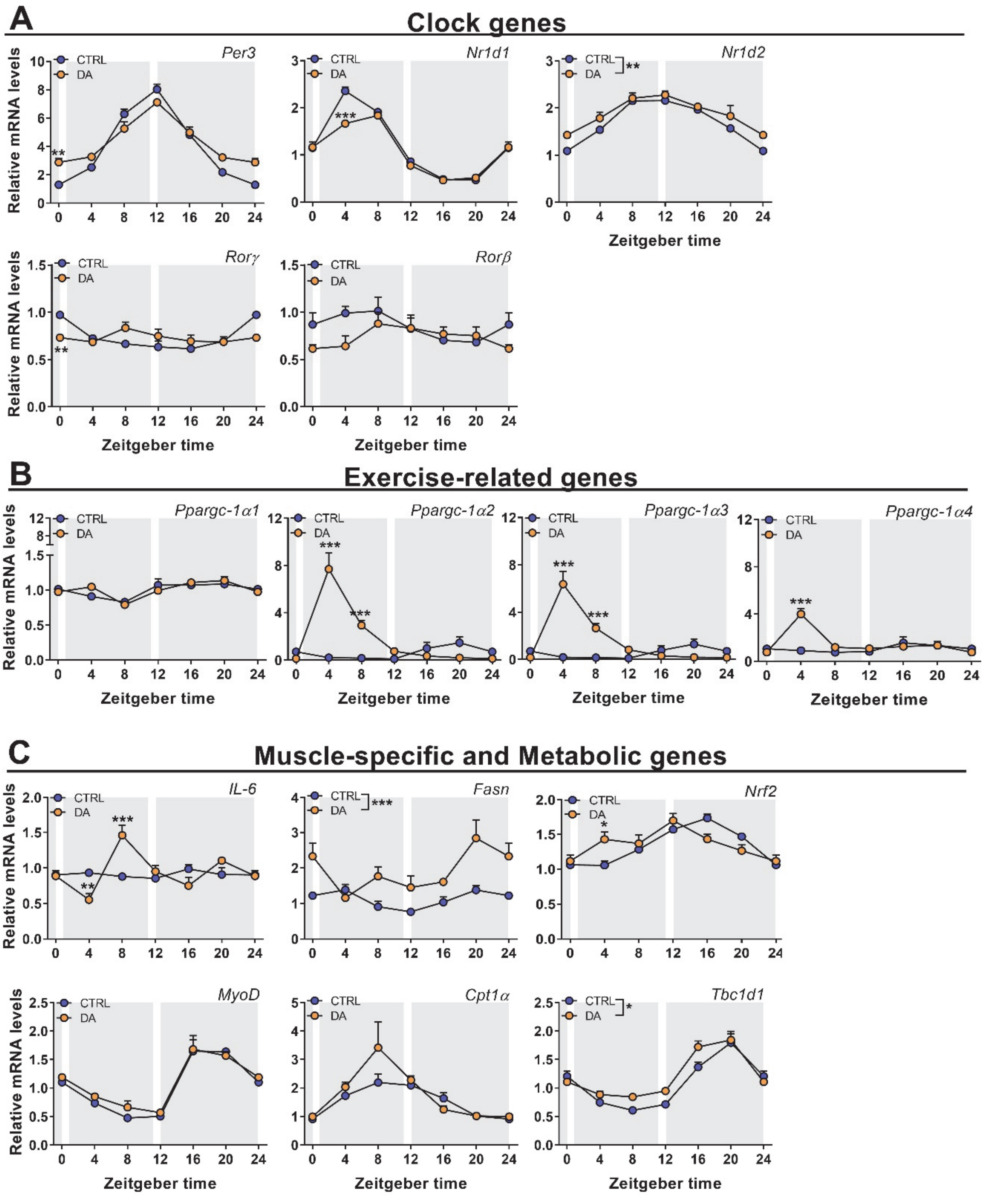


**SUPPL. FIGURE 5. Distinct muscle gene expression signatures after daytime vs. nighttime wheel running training.**

Gene expression in control (CTRL) and daytime activity (DA) mice. (A) Clock genes, (B) Exercise-related genes, (C) Muscle-specific and Metabolic genes. Expression values were determined by qPCR and normalized to *Hprt*. Data is shown as the average fold-change ± SEM (n = 4) relative to the expression in CTRL ZT0 set to 1 * $P < 0.05$; ** $P < 0.01$; *** $P < 0.001$. One-way ANOVA.

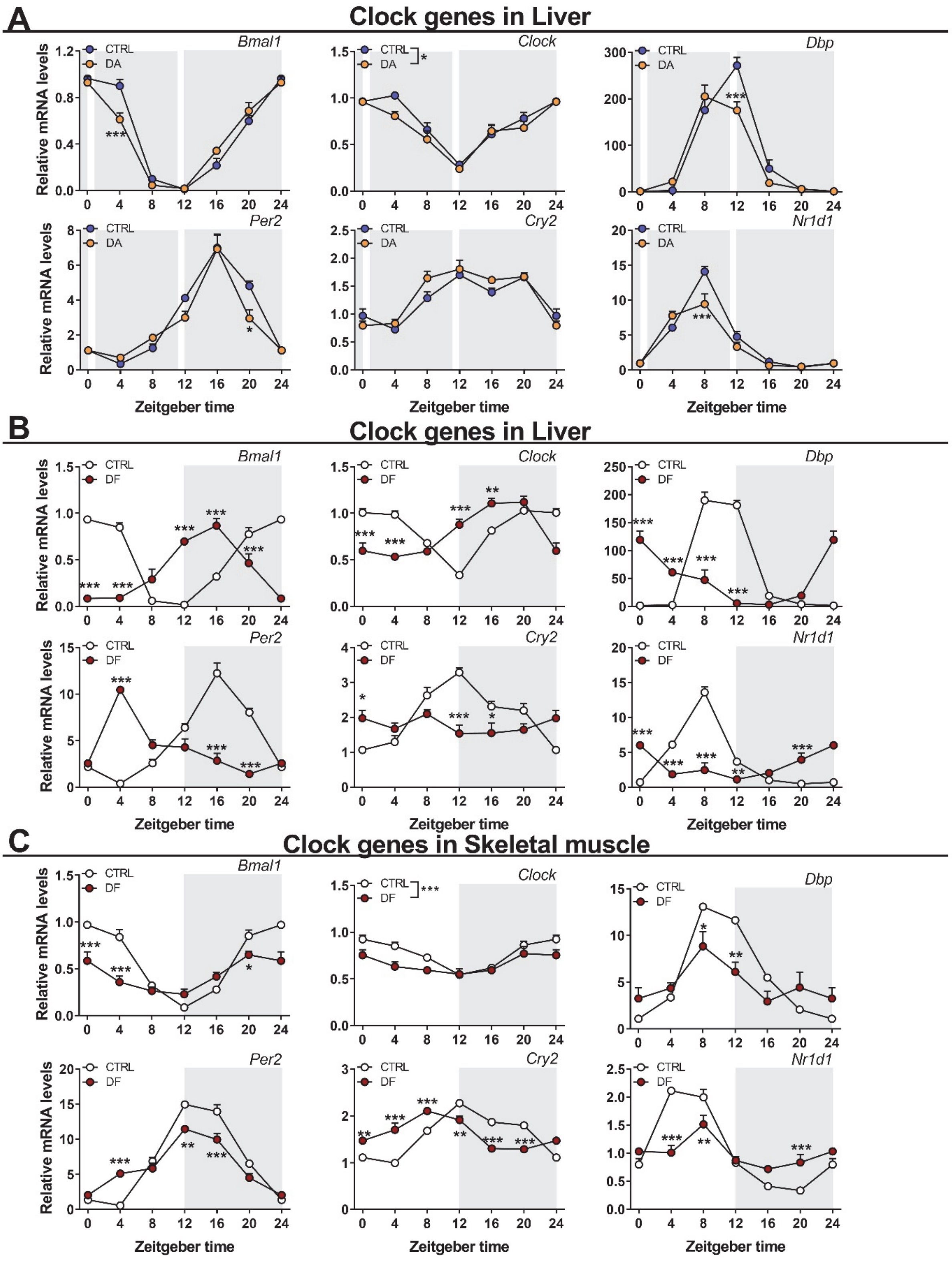


**SUPPL. FIGURE 6 Distinct liver clock gene expression signatures after daytime vs. nighttime wheel running training**

Gene expression in control (CTRL) and daytime activity (DA) mice. Clock genes in liver of DA mice (A), of daytime feeding (DF) mice (B), in skeletal muscle of DF mice (C). Expression values were determined by qPCR and normalized to *Hprt*. Data is shown as the average fold-change ± SEM ($n = 4$) relative to the expression in CTRL ZT0 set to 1 * $P < 0.05$; ** $P < 0.01$; *** $P < 0.001$. One-way ANOVA.

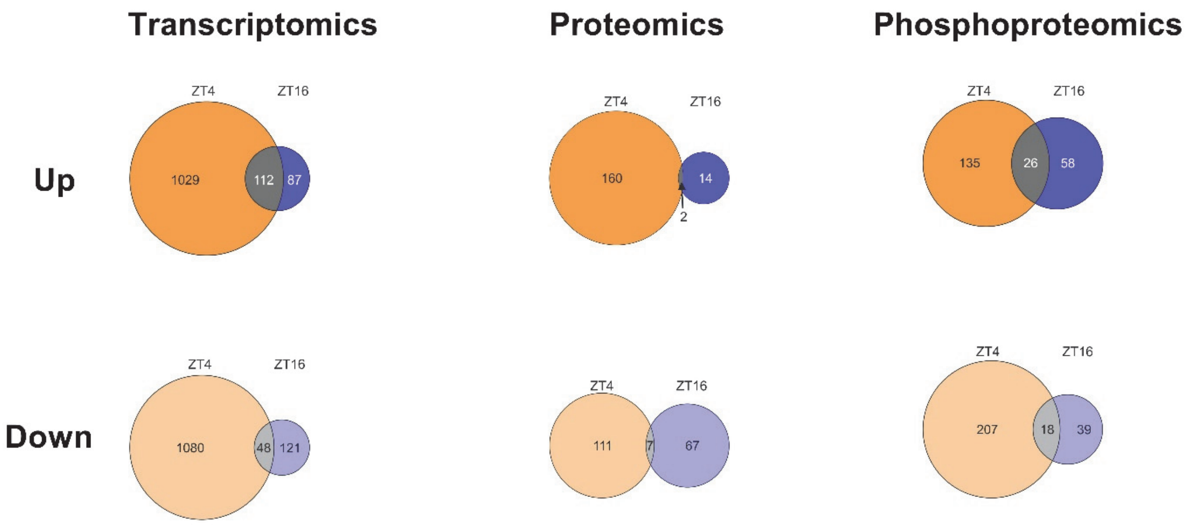


**SUPPL. FIGURE 7. Chronic voluntary daytime training influence the transcriptome, proteome, and phosphoproteome of skeletal muscle**

Venn diagram displaying the number of differentially regulated genes, proteins and phosphosites by nighttime (blue) or daytime (orange) wheel running, and the overlap (gray).
